# CRISPR antiphage defence mediated by the cyclic nucleotide-binding membrane protein Csx23

**DOI:** 10.1101/2023.11.24.568546

**Authors:** Sabine Grüschow, Stuart McQuarrie, Katrin Ackermann, Stephen McMahon, Bela E. Bode, Tracey M. Gloster, Malcolm F. White

## Abstract

CRISPR provides adaptive immunity in prokaryotes. Type III CRISPR systems detect invading RNA and activate the catalytic Cas10 subunit, which generates a range of nucleotide second messengers to signal infection. These molecules bind and activate a diverse range of effector proteins that provide immunity by degrading viral components and/or by disturbing key aspects of cellular metabolism to slow down viral replication. Here, we focus on the uncharacterised effector Csx23, which is widespread in *Vibrio cholerae*. Csx23 provides immunity against plasmids and phage when expressed in *Escherichia coli* along with its cognate type III CRISPR system. The Csx23 protein localises in the membrane using a N-terminal transmembrane α-helical domain and has a cytoplasmic C-terminal domain that binds cyclic tetra-adenylate (cA_4_), activating its defence function. Structural studies reveal a tetrameric structure with a novel fold that binds cA_4_ specifically. Using pulse EPR, we demonstrate that cA_4_ binding to the cytoplasmic domain of Csx23 results in a major perturbation of the transmembrane domain, consistent with the opening of a pore and/or disruption of membrane integrity. This work reveals a new class of cyclic nucleotide binding protein and provides key mechanistic detail on a membrane-associated CRISPR effector.

## Introduction

Type III (Csm and Cmr) CRISPR systems are multisubunit ribonucleoprotein complexes that are programmed by CRISPR RNA (crRNA) to detect invading RNA species [1]. Target RNA binding results in the activation of the catalytic Cas10 subunit, which has two possible active sites: a HD nuclease domain for ssDNA degradation [2–6] and a cyclase domain for generation of a cyclic oligonucleotide (cOA) signal [7, 8]. cOA molecules, which include cyclic tri-, tetra- and hexa-adenylate (cA_3_, cA_4_, cA_6_), act as messengers of infection, activating ancillary effector proteins. Characterised nuclease effectors include the Csm6/Csx1 family ribonucleases [7–12], the Can1 and Can2/Card1 nucleases and the exonuclease NucC [13–16]. The activation of these effectors can lead to non-specific degradation of key biomolecules, resulting in cell dormancy or programmed cell death [17, 18]. Type III CRISPR systems have been used as the basis for new diagnostic applications due to their ability to directly detect any desired RNA and generate an amplified cOA signal that in turn activates a reporter nuclease [19–21]. In some situations, cellular enzymes known as ring nucleases degrade the second messengers to revert cells to a non-infected ground state [22, 23]. Viruses also encode ring nucleases to subvert type III CRISPR immunity [24].

A wide range of putative type III CRISPR ancillary effector proteins have been predicted from bioinformatic studies [25, 26] and these are now being characterised, revealing new mechanisms for antiviral defence. For example, the cA_4_-binding protease CalpL has been shown to specifically degrade an anti-sigma factor, releasing an extra-cytoplasmic function (ECF)-family sigma factor to direct a transcriptional response to viral infection [27]. Many of the uncharacterised effectors appear to be membrane associated [25, 26]. For example, in the *Bacteroidetes*, an effector related to the magnesium transporter CorA can provide immunity when activated by a novel signalling molecule synthesised by the conjugation of ATP to S-adenosyl methionine [28].

Previously, we described a prophage-encoded type III-B (Cmr) CRISPR system from *Vibrio metoecus,* VmeCmr [21]. The VmeCmr Cas10 subunit lacks an HD nuclease domain and thus relies on cyclic nucleotide signalling for its function. On activation by cognate target RNA, VmeCmr generates predominantly cA_3_ on specific target RNA binding, resulting in the activation of the NucC effector nuclease for non-specific dsDNA degradation [16]. This phage-encoded system is a hybrid, using a type I-F Cas6 enzyme and associated CRISPR array for crRNA generation and a NucC or Csx23 effector [29]. Here, we demonstrate that Csx23 is a tetrameric membrane protein with a novel cytoplasmic cA_4_ recognition domain. Csx23 is activated by cA_4_ to prevent successful phage infection, most likely by disruption of the host membrane integrity.

## Results

### Csx23 functions as a type III CRISPR immune effector *in vivo*

Some *Vibrio* genomes host prophage-encoded type III CRISPR systems with an uncharacterised gene known as *csx23* (Figure 1A) [29]. The Csx23 family of proteins are commonly found in *Vibrio* species, where they appear as an alternative to NucC, and have a sporadic distribution associated with type III CRISPR systems in other bacterial phyla [26] (Figure S1).

**Figure 1.**
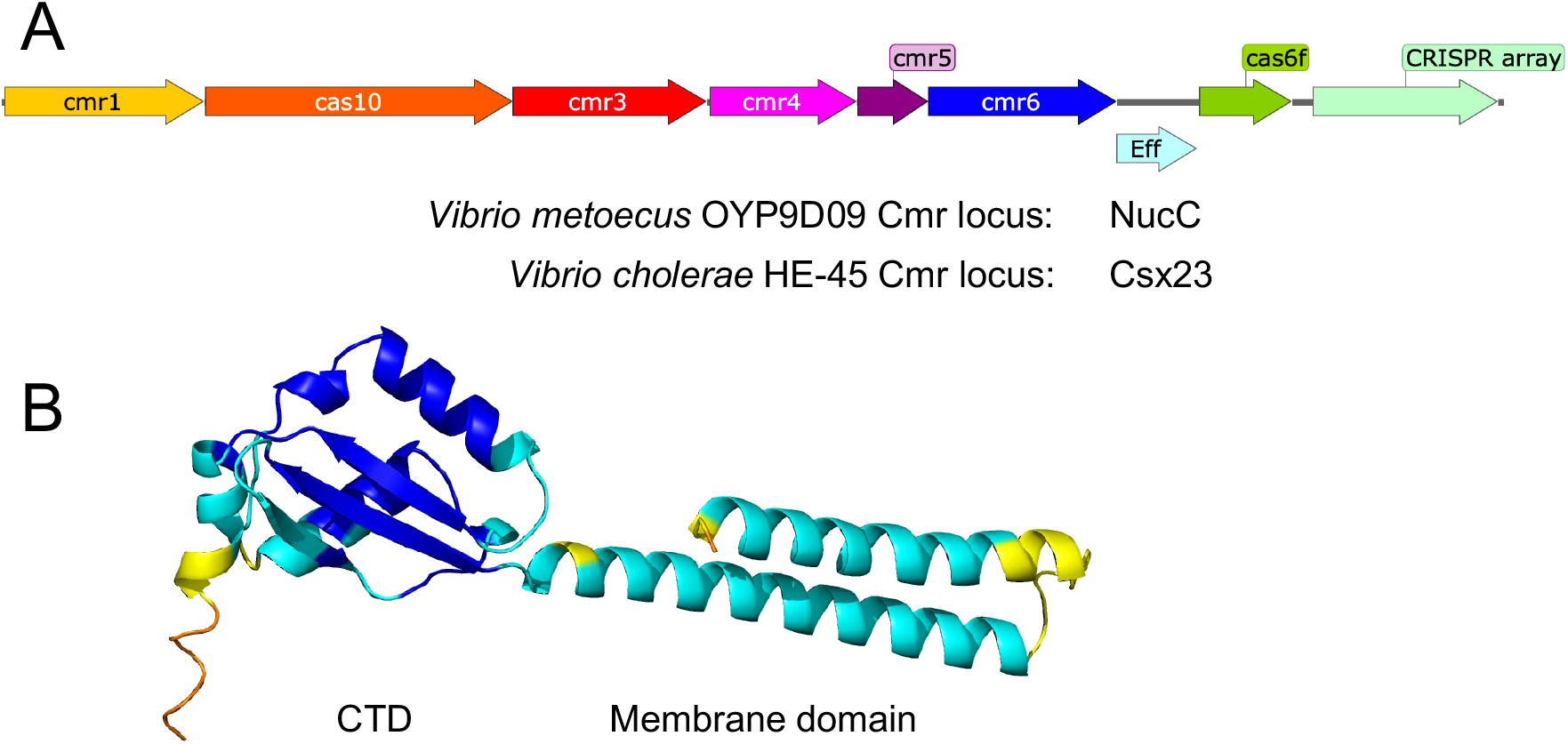
**A:** Organisation of *Vibrio* type III-B CRISPR (Cmr) loci. **B:** Predicted structure of *V. cholerae* HE-45 Csx23 (AF2) coloured by local distance difference test (LDDT) scaled from blue (high prediction confidence) to red (low).

We used AlphaFold 2 (AF2) [30], implemented on the Colabfold server [31], to predict the structure of Csx23 from *V. cholerae* HE-45. This yielded a model with a 68 residue N-terminal α-helical domain, predicted by InterPro [32] to be membrane spanning, and a 91 residue C-terminal domain (CTD) predicted to be a soluble, cytoplasmic domain (Figure 1B). The AF2 prediction was less confident for the arrangement of the domains relative to each other (Figure S2). We reasoned that the CTD would likely be the cOA-binding domain.

As a first step to elucidate whether Csx23 was indeed a cOA-dependent effector involved in adaptive immunity, we tested its activity in a plasmid challenge assay, making use of our established VmeCmr expression system [21]. In the assay, cells expressing the VmeCmr complex are transformed with a plasmid carrying an effector gene, *csx23* or *nucC*, alongside a target sequence for activation of the Cas10 cyclase. We used a portion of the tetracycline resistance gene as the target sequence and selected for tetracycline resistance after transformation of the cells with the target/effector plasmid. For active effectors, we expected to see fewer transformants compared to inactive effectors due to target depletion and/or programmed cell death (Figure 2A) [33].

**Figure 2.**
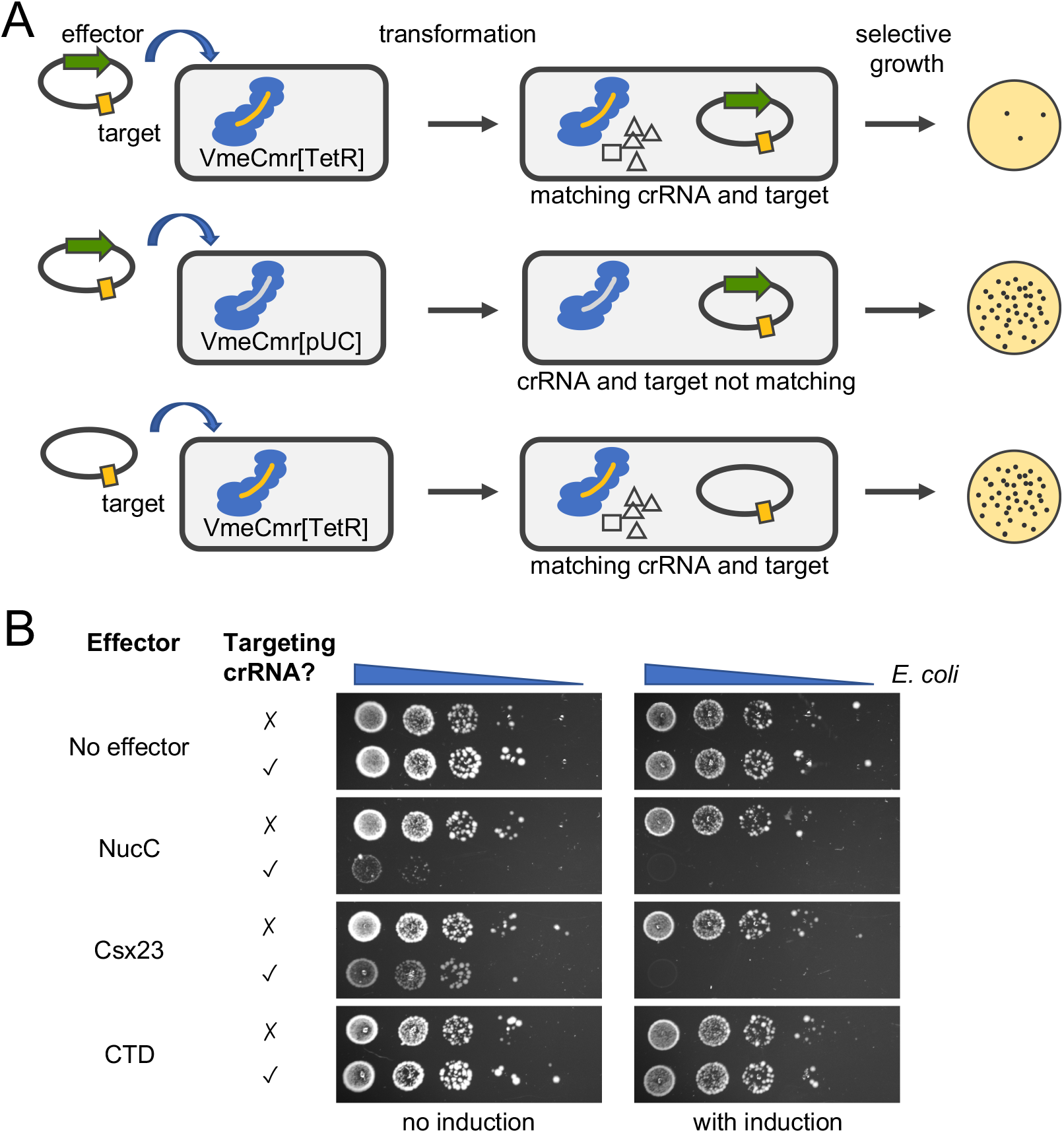
Plasmid challenge assay. **A:** Schematic representation of the basis for the assay. **B:** A serial dilution of the transformation mixture of *E. coli* cells expressing the VmeCmr complex with plasmids carrying a target sequence and varying effector genes was spotted onto agar plates containing antibiotics to select for all plasmids. VmeCmr complexes loaded with crRNA targeting the tetracycline resistance gene of the incoming plasmid (targeting crRNA, VmeCmr[TetR]) or loaded with crRNA targeting a sequence that is not present in the host genome or plasmids (non-targeting crRNA, VmeCmr[pUC]) were used.

In the absence of an effector gene, the same number of transformants was observed in the presence or absence of targeting crRNA, indicating that the VmeCmr system on its own does not confer any protection against invading nucleic acid, for example by Cas7-mediated target RNA knockdown (Figure 2B). In the presence of either NucC or Csx23, however, the number of transformants was significantly reduced in the presence of targeting crRNA. Using the AF2 model as a guide, we removed the membrane domain of Csx23 to only leave the CTD. CTD Csx23 was inactive in the plasmid challenge assay. These data confirmed that Csx23 can function as a Cmr-linked effector *in vivo*, that Csx23 function is dependent upon the presence of target and hence likely to be cOA-dependent, and that its membrane domain is essential for activity.

### Csx23 is a cA_4_-dependent, oligomeric effector

VmeCmr generates predominantly cA_3_ with only a minor amount of cA_4_ *in vitro*, and its effector nuclease NucC is activated by cA_3_ [21]. As the Csx23-associated VchCas10 was 99% identical at the amino acid level to VmeCas10, we reasoned that Csx23 would be activated by the same cOA species as NucC. To investigate cOA-binding by Csx23, we expressed and purified two versions of the protein: a full-length (FL) version, which required purification in the presence of detergent, and a truncated version encoding only the soluble CTD. The latter purified as a monomer of around 12 kDa (calculated MW 10 kDa) as determined by dynamic light scattering (DLS, Figure S3). FL Csx23 was isolated as a much larger entity of around 150 kDa (calculated MW 18 kDa for monomer). Membrane proteins purified in the presence of detergent form lipid-protein conglomerates which pose problems for standard methods to determine the oligomeric state of a protein such as analytical gel filtration or DLS. We therefore explored the quaternary structure of the Csx23 protein by cross-linking with bis(sulfosuccinimidyl)suberate (BS3) followed by SDS-PAGE analysis. The FL Csx23 protein could be cross-linked to form dimers, trimers and tetramers (Figure 3A). At the highest concentration of BS3, the tetrameric species was by far the dominant species, consistent with FL Csx23 existing as a tetramer. In contrast, the CTD did not cross-link in solution, suggesting a monomeric composition (Figure 3A).

**Figure 3.**
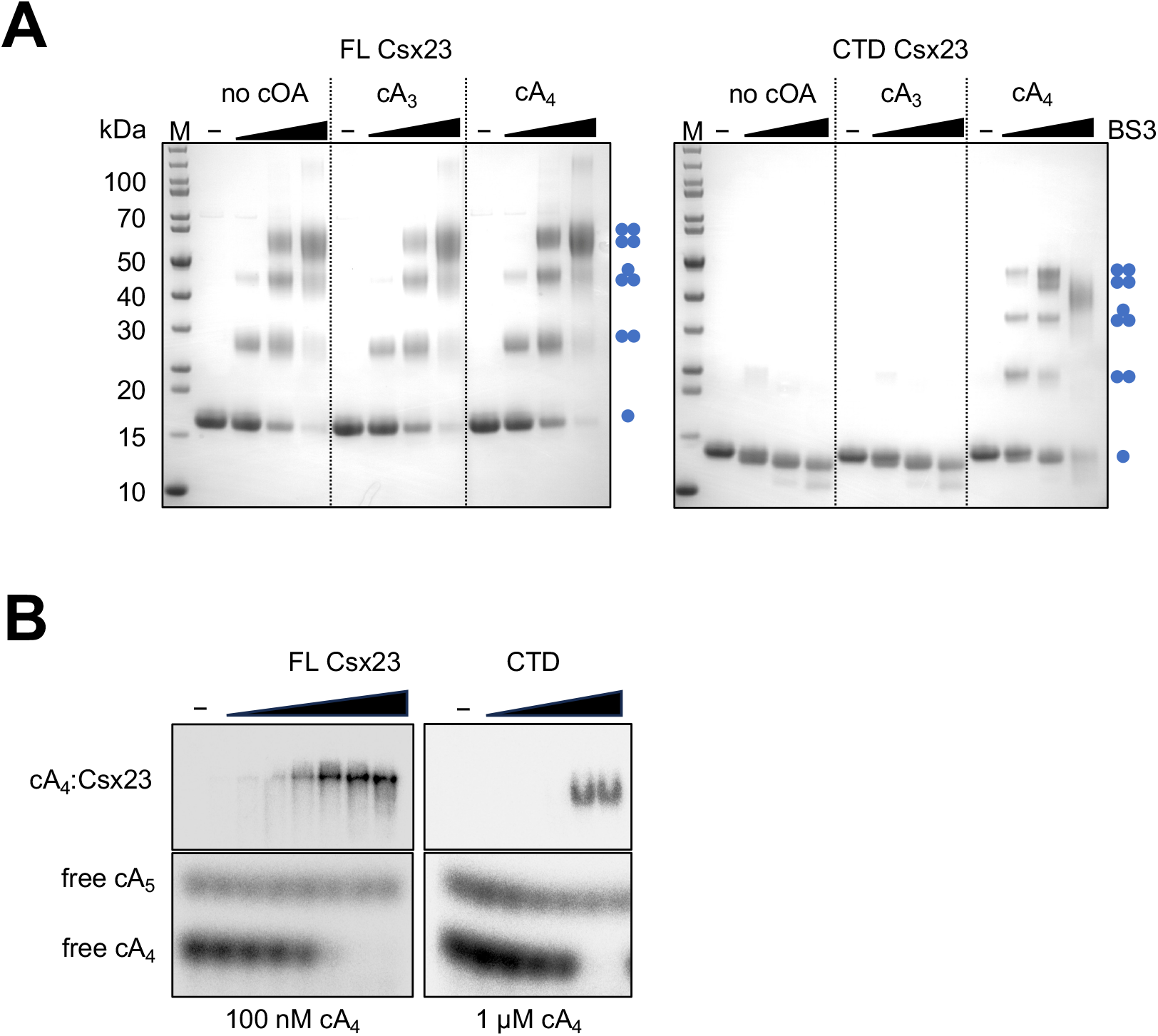
Csx23 is a tetrameric protein and binds cA_4_. **A:** BS3-Cross-linking of FL or CTD Csx23 in the presence and absence of cA_4_ or cA_3_. The FL protein formed tetramers in detergent regardless of cOA presence; the CTD could form tetramers in solution only in the presence of cA_4_. Blue dots show the predicted quaternary structure corresponding to each band on the SDS-PAGE. **B:** EMSA showing binding of radioactive cA_4_ by the FL and CTD Csx23 proteins. Protein concentrations were 25, 50, 100, 200, 400, 800 and 1600 nM for FL Csx3 and 0.4, 1, 4, 10, 40 and 100 µM for CTD Csx23. The full-length, uncropped gels are shown in Figure S4B.

We investigated the cOA-binding specificity of Csx23 by electrophoretic mobility shift assay (EMSA) using radioactively labelled cOA produced by the *Sulfolobus solfataricus* Csm complex, which predominantly produces cA_4_ and a small amount of cA_5_ [11] or by the VmeCmr complex that produces mainly cA_3_ with a trace of cA_4_ [21]. Unexpectedly, both FL and CTD Csx23 bound to cA_4_ but not cA_3_ (Figure 3B, Figure S4A), which strongly suggested cA_4_ as the relevant activator for Csx23. Cross-linking of both FL and CTD Csx23 in the presence of cA_4_ shifted the oligomeric state in SDS-PAGE from monomer to tetramer (Figure 3A). A 10 times higher concentration of CTD Csx23 was required to observe cA_4_ binding compared to FL Csx23, suggestive of weaker binding affinity (Figure 3B).

Although the specificity of Csx23 for cA_4_ rather than cA_3_ was contrary to our initial assumptions, it is frequently observed that the physiologically relevant cOA is not the most abundant species observed *in vitro*. For example, the Csm6 ribonucleases of both *Streptococcus thermophilus* and *Mycobacterium tuberculosis* are activated by cA_6_, which is only a minor component of the cOA mix produced *in vitro* [7, 33]. Furthermore, the distribution of cOA species produced by type III systems can differ *in vivo* compared to *in vitro* [34].

### Crystal structure of the tetrameric Csx23 soluble domain bound to cA_4_

A range of CRISPR effector proteins bind cA_4_, but they tend to utilize a conserved CARF/SAVED/Csx3 domain [10, 12, 13, 15, 23], which is a member of the Rossman fold superfamily [35] or the unrelated Crn2/AcrIII-1 domain [24]. The cA_4_ binding CTD of Csx23 was predicted to be completely unrelated to either family and could thus represent a new class of cA_4_ recognition domain. To investigate cA_4_ binding by Csx23 at an atomic level, we co-crystallised the soluble CTD of Csx23 with cA_4_. Diffraction data were collected on the crystals to a resolution of 1.76 Å, and the structure was solved using molecular replacement with the monomeric AF2 model as the search model.

The asymmetric unit contains a monomer of CTD Csx23, comprising two beta-strands, linked via two alpha-helices to a third beta-strand, which together form a mixed anti-parallel/parallel beta-sheet, followed by a short alpha-helix at the C-terminus (Figure S5A). The crystal structure of the monomer is consistent with the AF2 prediction, with a RMSD of 0.9 Å. Interestingly, the 4-fold crystallographic symmetry creates a tetrameric arrangement of CTD Csx23 (Figure 4A-C; Figure S5B, C), which is consistent with the behaviour of the protein in the presence of cA_4_ observed by cross-linking. Electron density in the *F*_obs_-*F*_calc_ map clearly showed the presence of a molecule of cA_4_ bound to CTD Csx23. The cA_4_ molecule, and a coordinating sodium ion, are positioned at the centre of rotation for the 4-fold crystallographic symmetry, meaning there is effectively one adenylate moiety bound to the Csx23 CTD monomer in the asymmetric unit; cA_4_ and the sodium ion were modelled with 0.25 occupancy to account for this. Further discussion will be based on the tetrameric structure bound to the whole molecule of cA_4_ as, due to the crystallographic symmetry, all interactions between each Csx23 CTD monomer and adenylate in cA_4_ are, by definition, identical.

**Figure 4.**
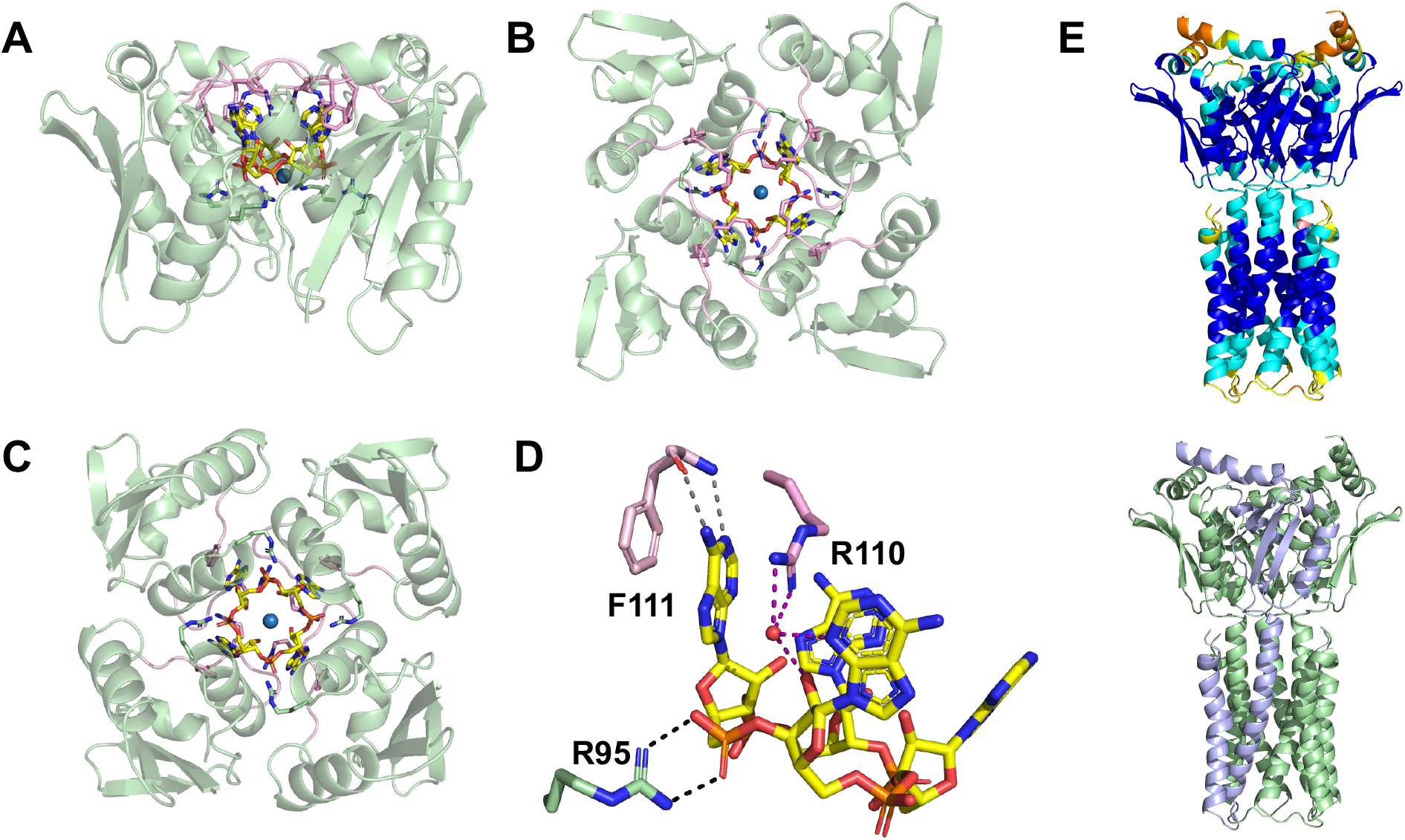
Structure of the tetrameric Csx23 CTD bound to cA_4_. **A:** Structure of tetrameric Csx23 CTD (green cartoon, with exception of flexible loop (residues 107-113) shown in pink) in complex with cA_4_ (sticks coloured by element, with carbon in yellow) from **A**: ‘side’, **B**: ‘top’, and **C**: ‘bottom’ views. The sodium ion is shown as a blue sphere. R95 (sticks coloured by element, with carbon in green) and R110 and F111 (sticks coloured by element, with carbon in pink) are also shown. **D**: Interactions formed between cA_4_ (yellow) and residues R95 (green), R110 and F111 (both pink) in Csx23 CTD. Colouring as in panels A-C. Residues from just one monomer are shown for clarity. Black dotted lines represent electrostatic interactions; grey dotted lines represent hydrogen bonds; purple dotted lines represent water-mediated hydrogen bonds. **E:** AF2 model of the FL Csx23 tetramer coloured by LDDT scaled from high (blue) to low (red) prediction confidence (top), and the same model with one subunit shown in light blue and the other three in light green (bottom).

The cA_4_ molecule is enclosed within the Csx23 CTD tetramer (Figure 4A-C; Figure S5D, E), but surprisingly makes few direct interactions with the protein. R95 of Csx23 CTD makes electrostatic interactions between both terminal nitrogen atoms and two different oxygen atoms in the phosphate group of cA_4_ (Figure 4D). R110 and F111 sandwich the adenine base in cA_4_ through ρε-ρε interactions. R110 also forms hydrogen bonds between both terminal nitrogen atoms and a water molecule, which in turn mediates hydrogen bonds with the C2-hydroxyl group of the ribose and a nitrogen atom in the adjacent adenine. Both the backbone amide and carbonyl groups of F111 form hydrogen bonds with different nitrogen atoms in the adenine base. A single sodium ion is positioned at the centre of the cA_4_, which interacts with a water molecule, but appears to make no direct or indirect interactions with cA_4_ or Csx23 CTD. Whilst R95 is positioned deep into the cA_4_ binding cavity, both R110 and F111 are located on a loop (comprising residues 107-113) on the surface of the Csx23 CTD (Figure 4A-C). This loop must display flexibility to allow cA_4_ access to the binding cavity, and then changes conformation to close over cA_4_ once bound. It is therefore likely that R110 and F111 are crucial for ‘locking’ the cA_4_ into position. The dynamic movement of loops has been observed previously with other structurally distinct domains that bind cOA [10, 14, 24, 36, 37]. These conformational changes are often accompanied by movement in other parts of the protein to elicit allosteric regulation. We used AF2 to model the tetrameric structure of FL Csx23 including the membrane spanning domain. This generated a model with high LDDT scores and predicted a transmembrane domain consisting of 8 α-helices (Figure 4E).

The cA_4_ forms a ‘cup-like’ structure, with the phosphodiester backbone forming the base of the ‘cup’ which is located deep in the binding cavity of Csx23 CTD (Figure S5F). The adenine bases are close to perpendicular to the backbone, thus forming the sides of the ‘cup’, and ensuring they are in a position near to the surface and thus accessible to residues R110 and F111 in the loop upon closing. The ribose moieties facilitate the formation of this distorted, and presumably high energy, conformation of cA_4_, by adopting an unusual ^2^_3_*T* twist conformation with C2’-*endo*/C3’-*exo* pucker. The conformation of cA_4_ bound to Csx23 CTD is distinct to that observed in complex with other cA_4_ binding proteins such as AcrIII-1 [24], Crn3 [23], Can1 [13], Can2/Card1 [14, 15] and Csx1 [10]. The angle between each C2’-hydroxyl group on the ribose and adjacent oxygen and phosphate atoms in cA_4_ is 156°. An angle close to 180° between these atoms is required for in-line nucleophilic attack to break the phosphodiester bond. Therefore, it is unlikely that Csx23 could support substrate-assisted ring nuclease activity, which is a feature of some self-limiting Csm6 family ribonucleases [12, 34, 36, 38]. Consistent with this, we observed no evidence for Csx23 ring nuclease activity *in vitro*.

The structural fold observed for Csx23 CTD is novel compared to other structures reported to bind cA_4_ (and other cOAs), demonstrating both the structural and functional diversity in the proteins that have evolved to interact with cyclic nucleotides. A DALI search [39] to identify any structural homologues of Csx23 CTD revealed intriguing matches to proteins containing PB1 and ubiquitin-like domains. The top hit, the PB1 domain of protein kinase C zeta type from rat [40], shares just 6% sequence identity with Csx23 CTD, but structurally overlaps with an RMSD of 3.1 Å over 66 alpha-carbon atoms (Csx23 CTD comprises 88 residues; PDB 4MJS; Figure S6A, B).

Superimposition of the two structures (Figure S6C, D) demonstrates a strong likeness between the secondary structure elements in the proteins, with just an additional beta-strand and loop present in the PB1 domain that is absent in Csx23 CTD. PB1 domains have a ubiquitin-like beta-grasp fold and are involved in protein-protein interactions in a host of biological processes [41]. PB1 domains have not previously been reported to exist in prokaryotes or viruses and the relevance of this structural relationship is unclear at present.

### Conserved residues important for cA_4_ binding

Based on the crystal structure, R95, R110 and F111 were implicated in cA_4_ binding. R95 and R110 are conserved in Csx23 homologues; F111 is well conserved but sometimes replaced by tyrosine, which can facilitate the same π-π interactions with the adenine base of cA_4_ (Figure S7). FL *csx23* mutants were first screened in the plasmid challenge assay (Figure 5A). Surprisingly, the F111A mutant showed wild type activity, but the R95A and R110A mutants allowed plasmid transformation, consistent with inactivation of Csx23. To explore this further, we purified FL Csx23 R95A and R110A and tested their ability to bind cOA species (Figure 5B). As expected, neither variant bound cA_4_, although Csx23 R95A unexpectedly showed a weak affinity for cA_5_.

**Figure 5.**
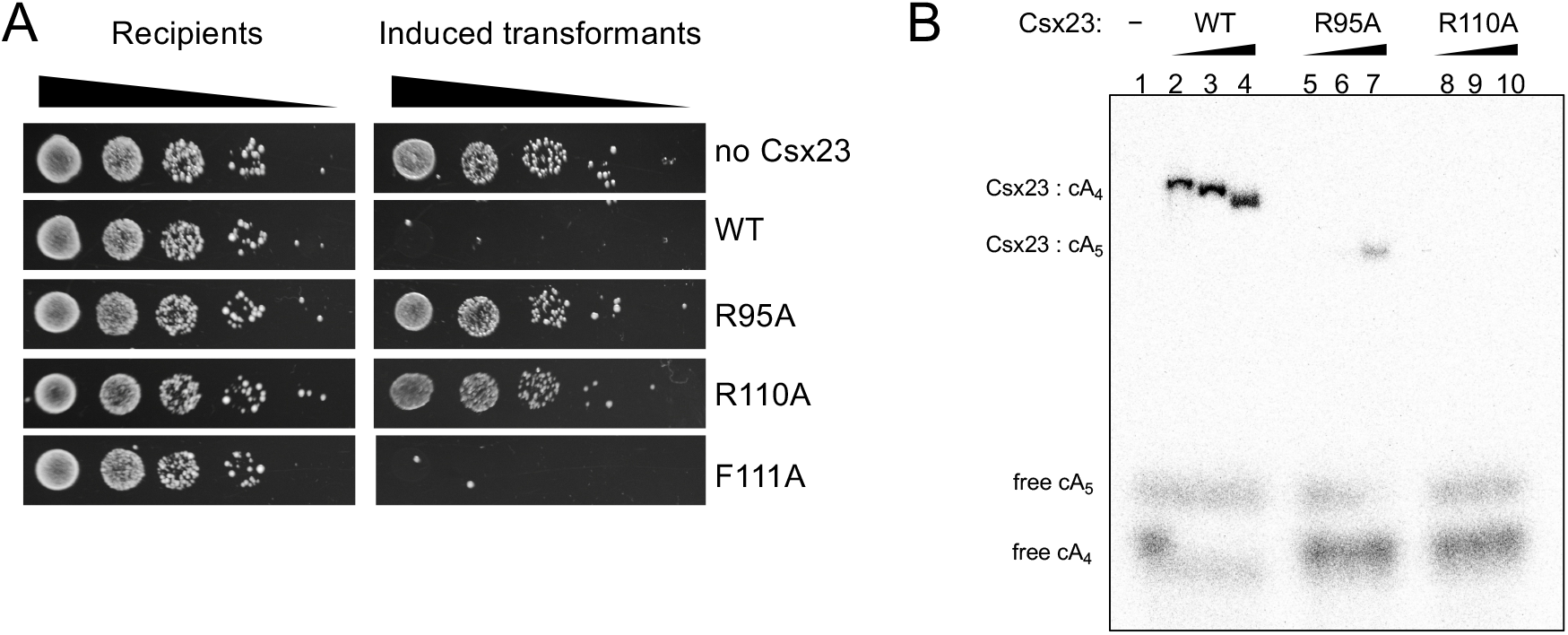
Investigation of roles of conserved residues. **A:** Plasmid challenge assay for wild-type and variant Csx23 proteins. The R95A and R110A variants did not provide immunity, suggesting Csx23 function was disrupted. **B:** EMSA showing that the R95A and R110A variants are unable to bind the cA_4_ activator *in vitro*. Weak binding of cA_5_ was observed for the R95A variant. Each binding reaction contained a two-fold molar excess of Csx23 tetramer over cOA. The amount of [^32^P]-cOA was kept constant, unlabelled cA_4_ was added to give final cA_4_ concentrations of 0.25 µM (lanes 2, 5, 8), 2.5 µM (lanes, 3, 6, 9), and 25 µM (lanes, 1, 4, 7, 10).

### Pulse EPR confirms the predicted transmembrane domain and its perturbation upon cA_4_ binding

Our working assumption for the activation of Csx23 was that binding of cA_4_ in the CTD results in a change in conformation that is communicated to the membrane spanning domain. Structural studies of cOA binding CARF family effectors have revealed that a “tightening” of the CARF domains around the bound ligand causes allosteric changes in the associated effector domains [14, 36, 37] and a similar scenario can be postulated for Csx23. As crystallisation of FL Csx23 was unsuccessful, we turned to Pulse Dipolar Electron Paramagnetic Resonance Spectroscopy (PDS) to obtain information about any conformational changes in the membrane spanning domain of Csx23 upon cA_4_ binding. PDS requires site-specific labelling of the protein of interest with a spin-label (to yield the paramagnetic side-chain R1); we chose to introduce cysteine residues at key locations for MTSL spin-label attachment. Wild type Csx23 contains 4 native Cys residues that were removed by primer-directed mutagenesis to give Csx23 AALA. Individual Cys residues were then introduced into the alpha-helices in the membrane domain to give Csx23 AALA V52C, N59C, or N62C. All variants were tested for activity using the plasmid challenge assay and showed similar results to the WT protein (Figure S8A). Subsequently the recombinant variants were purified in detergent (Figure S8B) and spin-labelled; labelling efficiency was determined by CW-EPR (Figure S9).

The PDS data showed the presence of two distances, as expected for a rotationally symmetric homotetramer. For the Csx23 AALA V52R1 (R1 refers to the spin label) variant, two sharp and narrow distance peaks were predicted by modelling the spin label rotamers both based on energy weighted rotamer modelling in MMM [42] and on excluded volume in mtsslWizard [43] using the AF2 predicted tetramer structure of Csx23. Experimental distance distributions were also obtained for this variant following reconstitution into nanodiscs, which agreed even better with the modelled distributions compared to the variant in detergent (Figure S10). The PDS data of the nanodisc reconstituted protein in the presence or absence of cA_4_ for Csx23 AALA V52R1 are shown in Figure 6. Notably, a dramatic change in distance distributions could be observed upon addition of cA_4_, suggesting a strong effect of the cyclic nucleotide on Csx23 conformation, with a substantially broadened distribution suggesting the sampling of a larger conformational ensemble. Similar results were obtained for the other two variants, with a broadening/shortening of the overall distribution for the N59R1 construct and a broadening leading to the loss of the resolution of the bimodality of the distance distribution for N62R1 (Figures S11 and S12). These observations indicate an increased conformational flexibility upon cA_4_ binding leading to structural heterogeneity in the transmembrane region.

**Figure 6.**
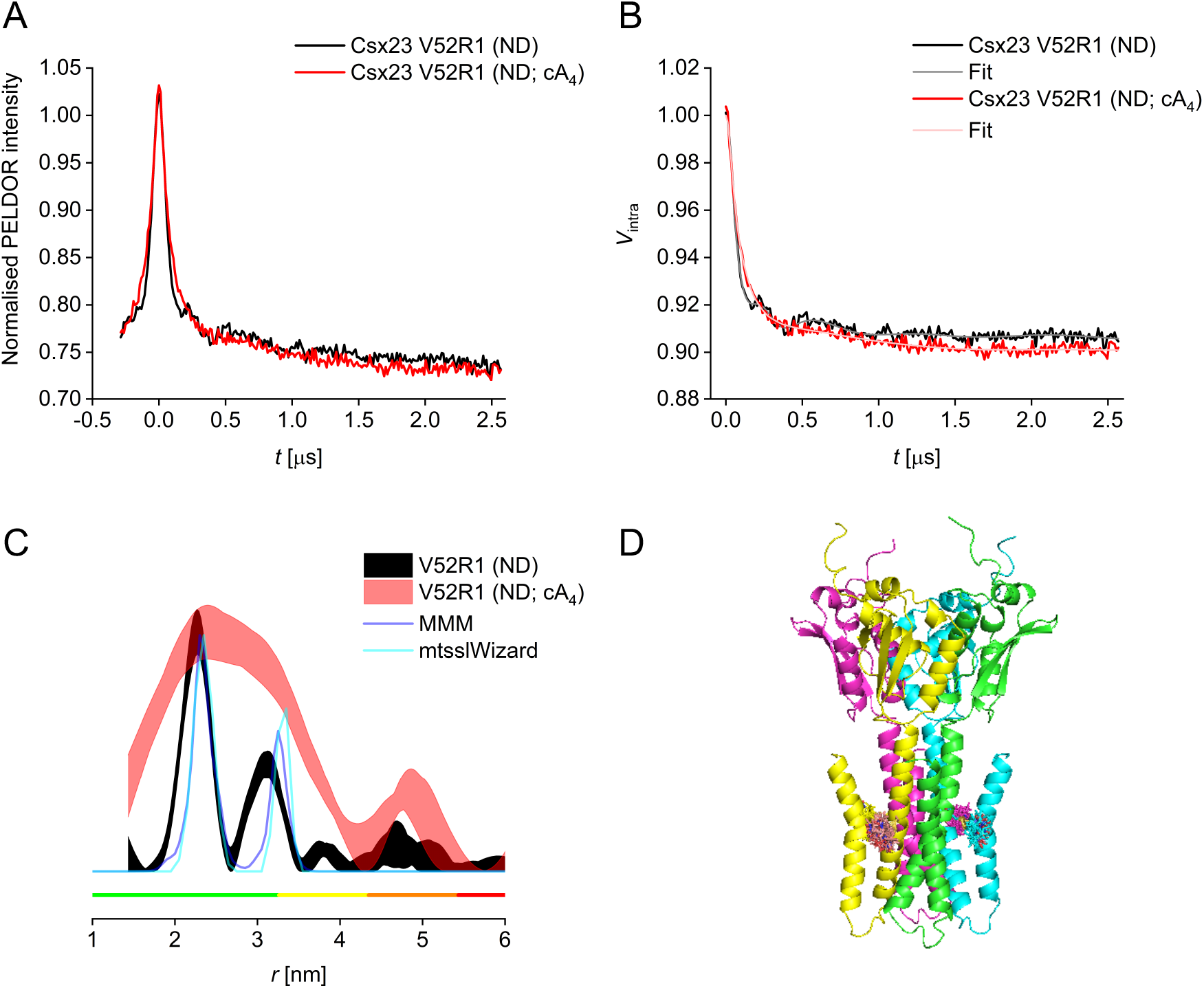
Pulse EPR demonstrates cA_4_ mediated perturbation of the transmembrane domain. Pulse electron-electron double resonance (PELDOR) data of the Csx23 AALA V52R1 variant reconstituted in nanodiscs (ND) in the presence (red) or absence (black) of cA_4_. **A:** Raw PELDOR data. **B:** Background-corrected traces with fits. **C:** Overlay of corresponding distance distributions shown as 95% confidence bands with predicted distributions from MMM and mtsslWizard based on the AF2 predicted tetrameric structure; colour bars indicate reliability ranges (green: shape reliable; yellow: mean and width reliable; orange: mean reliable; red: no quantification possible). **D:** Cartoon representation of AF2 predicted tetrameric structure of the spin-labelled Csx23 V52R1 variant; each monomer is shown in a different colour.

### Csx23 and NucC confer immunity to phage infection in *E. coli*

A limitation of the plasmid challenge assay is that it does not allow discrimination between a mechanism invoking effector-mediated programmed cell death and selective removal of the plasmid-encoded target, as both will lead to the same phenotype. To extend these studies, we proceeded to investigate the ability of the VmeCmr system to protect against infection by bacteriophage P1. *E. coli* cells expressing VmeCmr with a crRNA targeting the *lpa* gene of phage P1 [44] alongside the target- and effector-containing plasmid were infected with phage P1 at varying multiplicities of infection (MOIs) and the growth curves were recorded (Figure 7). As observed for the plasmid challenge assay, phage immunity by Csx23 and NucC was dependent on the presence of targeting crRNA and hence cOA production (Figure S13). In the absence of any effector, culture collapse occurred in an MOI-dependent manner 1 – 3 h post infection. The culture recovered approximately 10 h post infection (Figure 7A), either due to the establishment of stable prophages over time or due to the presence and subsequent proliferation of persister cells. However, when either Csx23 or NucC were expressed, the growth curves of cells infected with phage P1 at low to moderate MOIs were almost indistinguishable from those of uninfected cells, suggesting efficient anti-phage defence. At high MOI (15), where almost all cells should be infected by phage, culture collapse occurred approximately 1 h post infection in the absence of Csx23, as expected (Figure 7B). However, in the presence of Csx23 the cells responded in a markedly different way. Cell growth stopped earlier (30 vs 60 min), without a pronounced crash in OD_600_ and cell growth recommenced much more quickly than in the absence of effector. Similar behaviour was observed for cells expressing NucC (Figure S13). The early growth arrest observed here could fit with a programmed cell death (abortive infection) mechanism, as suggested previously for NucC [16], but the growth behaviour overall appeared more consistent with a mechanism involving cell dormancy rather than death.

**Figure 7.**
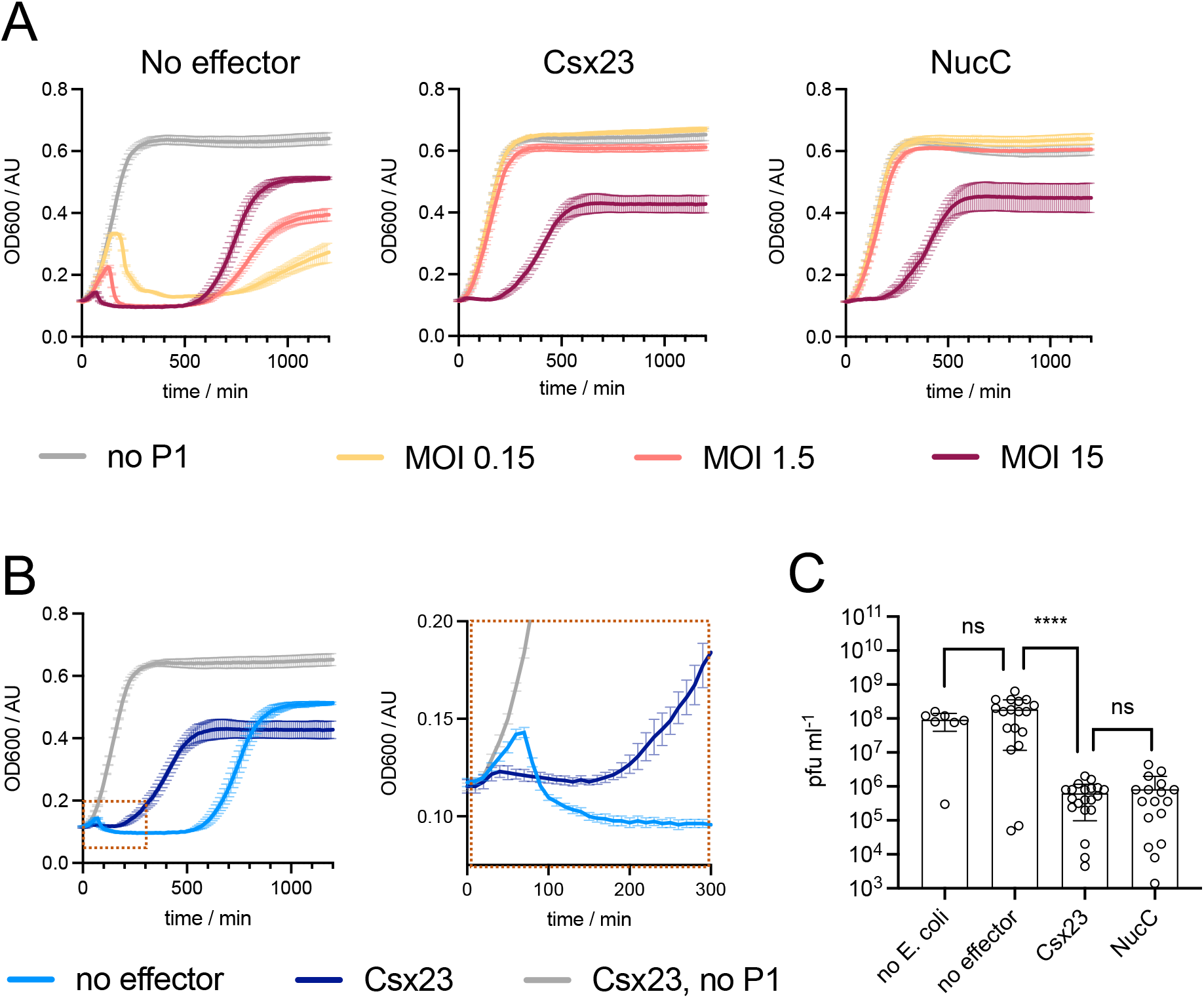
Phage P1 challenge of *E. coli* expressing VmeCmr and varying effector genes. **A:** Growth curves of cells with no effector, Csx23 or NucC with increasing amounts of phage P1. MOI: multiplicity of infection. The growth curve in the absence of phage infection is shown for comparison. **B:** Growth curves from cells carrying Csx23 or no effector after phage P1 infection at MOI 15 and expanded view to show early time points. **C:** The number of viable phage particles was determined by applying a dilution series of cleared supernatant from infected cultures to agar plates overlaid with BL21(DE3) Star cells, the same *E. coli* strain as used in all *in vivo* assays. The plaques from ≤ 3 independent experiments consisting of 2 biological replicates each were counted and plotted using Graphpad Prism. Mann-Whitney test (Prism 10.0) was used to determine statistical difference (not significant (ns): *p-*value ≤ 0.05, ****: *p*-value <0.0001). Cultures of strains containing either Csx23 or NucC contained 100 times fewer viable phage compared to those that did not carry any effector.

Irrespective of the presence of any effector protein in the host, the *E. coli* cultures grew to a similar density given enough time (> 18 h). We therefore tested whether there were any differences in the amount of viable phage P1 between the strains 24 h post infection. Strains containing phage P1-targeting VmeCmr expressed with Csx23, NucC or no effector were infected at an MOI of 15. A serial dilution of the cleared culture supernatant 24 h post infection was then applied to agar plates containing an indicator strain and the phage titre was determined from the number of plaques after overnight incubation (Figure 7C). In the presence of either NucC or Csx23, a reduction in viable phage particles of at least 2 orders of magnitude was observed compared to cultures without any effector, confirming interference of phage P1 proliferation. The significance of these observations is discussed below.

## Discussion

Type III CRISPR systems utilize a wide range of cyclic nucleotide-activated effector proteins for defence against mobile genetic elements [25, 26]. The system studied here is encoded on a prophage that is found in many *V. cholerae* and *V. meteocus* genomes [29]. As such, it may function in inter-MGE conflict within the host cell, rather than providing anti-phage defence *per se*. Previously, we showed that this system remains activated for an extended period, generates large quantities of cOA and is associated with the robust endonuclease effector NucC [21] – all features that may reflect the priorities of the prophage to prevent superinfection by competitors, at the expense of the host cell if necessary.

Membrane-associated CRISPR effectors have been detected bioinformatically but characterised examples are still scarce. In addition to the aforementioned *B. fragilis* CorA effector [28], some type V CRISPR systems have an associated Csx28 protein that forms an octameric membrane pore, resulting in membrane depolarisation, although the mechanism of activation remains unclear [45]. More broadly, membrane associated effectors function in many other prokaryotic defence pathways that are presumed to function via programmed cell death. One example is the Aga2 protein, which depolarises membranes when an associated Argonaute protein is activated during viral infection in *Saccharolobus islandicus* [46]. Furthermore, the CBASS and Pycsar anti-phage defence systems, which also function via cyclic nucleotide signalling, frequently encode effectors that function via membrane disruption [47, 48]. To date, only the CBASS Cap15 effector has been investigated in detail. It is activated by binding a cyclic dinucleotide and oligomerizes to disrupt and depolarise cell membranes, leading to cell death [49].

Csx23 utilises a small tetrameric cytoplasmic domain to bind cA_4_ – representing a new class of cyclic nucleotide recognition domain. The crystal structure of the CTD of Csx23 revealed that mobile loops harbouring key interacting residues for cA_4_ enclose the cyclic nucleotide binding cavity, reminiscent of the CARF and Crn2 families [50]. EPR data show that cA_4_ binding to the CTD of Csx23 results in the structural perturbation of the trans-membrane helical domain, consistent with the hypothesis that Csx23 functions via membrane disruption, possibly as a type of ligand-gated channel. We observed effective Csx23-medited anti-phage immunity but not a classical abortive infection phenotype, as infection at a very high MOI did not lead to complete culture collapse. This should be caveated by the fact that we overexpressed the defence system in *E. coli*, and by the understanding that phage P1 is a temperate phage. Nonetheless, these data do not provide strong support for a simple model of immunity by altruistic suicide and are perhaps more consistent with the induction of dormancy.

In conclusion, our work reveals a new class of membrane-associated CRISPR effector that functions via cyclic oligoadenylate signalling to provide anti-viral immunity. The cA_4_ binding domain of Csx23 is unrelated to the near-ubiquitous CARF/SAVED domain, highlighting the potential for further discoveries in the mechanistically diverse type III CRISPR and CBASS defence pathways. Although we observe a cA_4_-activated allosteric modulation of the Csx23 trans-membrane domain structure we have not yet observed changes in membrane permeability – despite significant efforts to do so. This is a clear area for future study.

## Materials and Methods

### Sub-cloning and site-directed mutagenesis

Enzymes were purchased from Thermo Scientific or New England Biolabs and used according to manufacturer’s instructions. Oligonucleotides and synthetic genes were obtained from Integrated DNA Technologies (IDT, Coralville, Iowa, USA). Synthetic genes were codon-optimized for *E. coli* expression and restriction sites for cloning incorporated where necessary. All final constructs were verified by sequencing (GATC Biotech, Eurofins Genomics, DE). *Vibrio cholerae* HE-45 *csx23* (Csx23) was obtained as a G-Block with flanking restriction sites for cloning. After digestion with *Nco*I and *Bam*HI, *csx23* was ligated into linearised pEV5HisTEV [51]. The expression construct for full length Csx23 with a non-cleavable C-terminal His-tag was obtained by PCR-amplifying Csx23 from the above construct using primers 5’-ATGGCACATATGAATACTTTCAAGCG and 5’-atatctcgagTGCATTGGAGCCGCTCTTGG. After digestion with *Nde*I and *Xho*I, the gene was ligated into pEV5HisTEV. Expression constructs for the C-terminal domain of Csx23 (Csx23_CTD_) were obtained using the same strategy. For expression with an N-terminal His-tag, 5’-acttccatgGACGAGATCACTGTCGTCCTG and 5’-GGAGCTCGAATTCGGATCCCT were used as primers and the digested PCR product was inserted into *Nco*I / *Xho*I sites of pEV5HisTEV. For expression with a C-terminal His-tag, 5’-actctcatatgAATGACGAGATCACTGTCGTCCTG and 5’-atatctcgagTGCATTGGAGCCGCTCTTGG were used as primers and the digested PCR product was inserted into the *Nde*I / *Xho*I sites of pEV5HisTEV. Csx23 variants for protein isolation were obtained using primer-directed mutagenesis of the expression constructs described above. Expression constructs for EPR was obtained through successive rounds of primer-directed mutagenesis.

### Constructs for plasmid challenge and phage immunity assays

The VmeCmr1-6 expression construct (pVmeCmr1-6) contained a ColE1 origin of replication and ampicillin resistance gene, and the expression of *cmr1-3* and *cmr4-6* was driven by individual T7 promoters. The *cmr2* (*cas10*) gene included a sequence encoding a TEV-cleavable, N-terminal His_8_-tag. The construction of this plasmid has been previously described [21]. Production of crRNA was achieved by placing a minimal *V. metoecus* CRISPR array and *V. metoecus cas6f* into pCDFDuet™-1 (Novagen, Merck Millipore) as previously described [21]. The CRISPR array contained two repeat sequences flanking two oppositely directed BpiI recognition sites (5’-gtgtcttcgtaccttgaagacca) to allow later insertion of the target/spacer sequence of choice. Spacer sequences were obtained as synthetic oligonucleotides with a 5’-overhang sequence of 5’-GAAA for the sense strand and 5’-GAAC for the antisense strand. After the two strands were annealed, they were ligated into BpiI-digested pCDFDuet-derivative. vmeRepeat and spacer sequences are listed in Table S1. The pCDF derivatives are named according to the gene targeted by the spacer sequence as pCRISPR^Target^.

Effector proteins were cloned into pRATDuet [33]. This plasmid carries the pRSF1030 replicon from RSFDuet™ (Novagen, Merck Millipore), *araC* and the *araBAD* promoter from pBAD/His (Invitrogen), the tetracycline resistance gene from pACE2 (Geneva Biotech, Genève, CH) and the two MCSs from pACYCDuet™-1 (Novagen, Merck Millipore). *Csx23* was inserted into MCS-1 of pRATDuet (NcoI/SalI) from the *Nco*I / *Xho*I-digested gBlock™. *NucC* was inserted into MCS-2 of pRATDuet (*Nde*I/*Xho*I) from the *Nde*I / *Xho*I-digested gBlock™ and subsequently two nucleotides between the RBS and translational start codon were removed by primer-directed mutagenesis to reduce the amount of NucC being produced.

Csx23 variants used in plasmid challenge assays were constructed by single or successive rounds of primer-directed mutagenesis using the pRATDuet construct as template and the same primers as used for mutagenesis of expression constructs.

### Protein Production and Purification

*E. coli* C43(DE3) was used as the expression host for full length Csx23 and Csx23_CTD_. Overnight cultures were diluted 100-fold into LB containing 50 µg ml^-1^ kanamycin, and incubated at 37 °C, 220 rpm until the OD_600_ reached 0.6 – 0.8. After induction with 200 µM IPTG, incubation was continued at 37 °C for 4 h. Cells were harvested by centrifugation and pellets stored at -20 °C. For Csx23_CTD_, cells were resuspended in lysis buffer (50 mM Tris-HCl, 500 mM NaCl, 20 mM imidazole, 10% glycerol, pH 7.6) and lysed by sonication. The lysate was cleared by ultracentrifugation (40,000 rpm, 40 min, 4 °C) and loaded onto a HisTrap FF 5 ml column (Cytiva), washed with 5 CVs lysis buffer, then with 6 CVs 4% elution buffer (50 mM Tris, 0.5 M NaCl, 0.5 M imidazole, 10% glycerol, pH 7.6) and protein was eluted in a gradient over 10 CVs leading to 100% elution buffer. Csx23_CTD_-containing fractions were dialysed at room temperature overnight in the presence of TEV protease against lysis buffer. The protein solution was passed through the HisTrapp FF column a second time, and the flow-through was concentrated using an Amicon Ultracentrifugal filter (3 kDa MWCO, Merck-Millipore) and further purified by gel filtration (HiLoad™ 16/600 Superdex™ 200 gp, Cytiva) using 20 mM Tris-HCl, 250 mM NaCl, 10% glycerol, pH 7.5 as mobile phase. Csx23_CTD_-containing fractions were pooled and concentrated.

For full length Csx23, cells were resuspended in lysis buffer containing 1% (*w/v*) DDM (*n*-dodecyl-beta-maltoside) and lysed by sonication. After stirring at room temperature for 1 h, the lysate was cleared by ultracentrifugation (40,000 rpm, 40 min, 4 °C) and loaded onto a HisTrap FF 5 ml column (Cytiva), washed with 5 CVs lysis buffer containing 0.1% DDM, then with 7 CVs 4% elution buffer containing 0.1% DDM. The protein was eluted in a 10 CV gradient to 100% elution buffer containing 0.1% DDM. Csx23-containing fractions were concentrated using an Amicon Ultracentrifugal filter (30 kDa MWCO, Merck-Millipore) and further purified by gel filtration (HiLoad™ 16/600 Superdex™ 200 gp, Cytiva) using 20 mM Tris-HCl, 250 mM NaCl, 10% glycerol, 0.06% DDM, pH 7.5 as mobile phase. Csx23-containing fractions were pooled and concentrated. Csx23 variants were purified as described for wild type Csx23.

The proteins were stored at 4 °C for short term storage or flash-frozen as single-use aliquots and stored at -80 °C. The concentration of full length Csx23 and Csx23_CTD_ was determined spectrophotometrically by absorbance at 280 nm and using the calculated extinction coefficients of 9970 and 1490 M^-1^ cm^-1^, respectively. Concentrations are stated for the monomer.

### Dynamic Light Scattering

DLS measurements were performed with the Zetasizer Nano S90 (Malvern) instrument. Samples contained 60 – 100 µM protein in 20 mM Tris-HCl, 75 mM NaCl, 10 mM MgCl_2_ and 0.1% DDM as required. After centrifugation at 12,000 x g for 10 min at room temperature, the sample was filtered (0.22 μm PES membrane) and loaded into a quartz cuvette (ZMV1012). Measurements were performed at 25 °C with three measurements of thirteen runs.

### Electrophoretic Mobility Shift Assay

[α-^32^P]-Radiolabelled cA_3_ and cA_4_ were prepared using the VmeCmr or SsoCsm complex, respectively, as previously described [21, 22]. A typical EMSA assay contained 50 nM [a-^32^P]-cA_4_ and 0 – 1600 nM Csx23 in 20 mM Tris, pH 8.0, 100 mM NaCl, 10% glycerol, 0.5 mM TCEP. The mixture was incubated at room temperature for 10 min before addition of G-250 Native Gel Sample Loading Buffer (Invitrogen) to 0.005% final concentration. Samples were immediately loaded onto a pre-run 6% acrylamide gel (29:1 acrylamide:*bis*-acrylamide). Radiolabelled material was separated for 2 h at 200 V in 1X TBE buffer and visualised by exposure to a phosphor storage screen (Cytiva).

### Protein Crosslinking

Csx23 was dialysed against 20 mM HEPES, 150 mM NaCl, 0.1% DDM if required, 10% glycerol, pH 7.6 for 2 h at room temperature. The reaction was carried out in dialysis buffer with 40 µM Csx23 in the absence or presence of 80 µM cOA and 0 – 3.3 mM BS-3 (bis(sulfosuccinimidyl)suberate-d_0_, Thermo Scientific) crosslinker and allowed to proceed with gentle agitation for 30 min at 20 °C. SDS-PAGE sample loading buffer was added to each reaction and the samples were heated to 95 °C for 2 min before analysis by SDS-PAGE.

### Plasmid Challenge Assay

*E. coli* BL21 Star™ (DE3) (Invitrogen) was transformed with pVmeCmr1-6 and pCRISPR^TetR^ (tetracycline resistance gene-targeting crRNA) or pCRISPR^pUC^ ^MCS^ (pUC19 MCS-targeting crRNA, non-targeting control). Competent cells were prepared from fresh individual transformants as follows: a culture obtained from 50 – 100-fold dilution of an overnight culture was grown at 37 °C, 220 rpm to an OD_600_ of 0.4 – 0.6. All subsequent steps were carried out at 4 °C with pre-chilled buffers. Cells were collected by centrifugation (10 min, 3000 rpm, Eppendorf centrifuge 5810) and the pellet resuspended in an equal volume of 60 mM CaCl_2_, 25 mM K-MES, pH 5.8, 5 mM MgCl_2_, 5 mM MnCl_2_. After incubation on ice for 1 h, cells were harvested again and resuspended in 1/10^th^ volume of the same buffer containing 10% glycerol. Single-use aliquots were stored at -70 °C. For the plasmid challenge assay, 50 ng (1 µl) target plasmid containing the effector was added to 50 µl of competent cells. After incubation on ice for 30 min, cells were heat-shocked at 42 °C for 1 min, then placed on ice for 2 min. After the addition of 0.5 ml LB medium, the transformation mixture was incubated for 2 – 2.5 h at 37 °C, 220 rpm. A 10-fold serial dilution (2.5 µl / spot) was applied onto selective LB agar plates and the plates were incubated at 37 °C for 20 h.

### Phage Immunity Assay

*E. coli* BL21 Star™ (DE3) (Invitrogen) was co-transformed with three plasmids: one encoding the VmeCmr complex (pVmeCmr1-6), one encoding the crRNA (pCRISPR^Lpa^ for phage P1 *lpa* gene-targeting crRNA or pCRISPR^pUC^ ^MCS^ as non-targeting control), and one encoding the effector (pRAT-Csx23, pRAT-NucC or pRATDuet with no insert). Single colonies were grown overnight in LB medium containing 50 µg ml^-1^ ampicillin, 25 µg ml^-1^ spectinomycin and 6.5 µg ml^-1^ tetracycline at 25 – 28 °C with shaking. The overnight cultures were either used for the phage immunity assay (see below) or aliquots (0.3 ml) were collected, mixed with 50% glycerol and stored at -70 °C for future use. Aliquots were revived by inoculating them into 5 ml LB containing the appropriate antibiotics followed by a 2 – 3 h outgrowth period at 37 °C with shaking.

The relationship between OD_600_ and cell density for BL21 Star carrying the three plasmids was determined by two independently performed microdilution assays. An OD_600_ of 1.0 yielded around 2·10^8^ cfu ml^-1^ for *E. coli* BL21 Star™ (DE3) carrying the three plasmids.

For the phage immunity assay, the OD_600_ for each culture was determined and each culture was diluted to an OD_600_ of 0.05 (∼1·10^7^ cfu ml^-1^) with LB medium supplemented with 10 mM MgSO_4_, 0.2% *w/v* l-arabinose and the appropriate antibiotics. It had been found that the presence or absence of antibiotics made no difference to the assay in the first ∼ 5 h, hence antibiotics were omitted for shorter time courses. 160 µl *E. coli* was added to each well (96-well propylene clear flat-bottom plate, Greiner) alongside supplemented LB without *E. coli* as control and background. Phage P1 was diluted from a >1·10^10^ pfu ml^-1^ stock into the supplemented LB broth and 40 µl was added to each well to give final MOIs quoted in the individual experiments. Growth curves were monitored by OD_600_ readings at 37 °C (FluoStar Omega plate reader, BMG Labtech). The absorbance at 600 nm was recorded in matrix scan mode (2×2 matrix, 2 mm scan width, 0.5 s settling time, 10 flashes per well and cycle). The plate was shaken for 10 s at 200 rpm in double orbital mode before each cycle with a typical cycle time of 15 min.

Phage titre was determined by microdilution drop assay of culture aliquots that were collected 24 h post infection and centrifuged for 3 min at 2260 x g. The supernatant was carefully transferred to a fresh tube, and a serial 10-fold dilution was prepared using LB containing 10 mM MgSO_4_. 2.5 µl phage dilution was applied onto soft agar containing 10 mM MgSO_4_ and *E. coli* BL21 Star™ (DE3) at OD_600_ 0.2 with a layer of LB agar/MgSO_4_ underneath. Plates were incubated at 37 °C overnight before counting plaques. Results were plotted using Graphpad Prism. Mann-Whitney test (Prism 10.0) was used to determine statistical significance.

### Sample preparation for Pulse Dipolar Electron Paramagnetic Resonance Spectroscopy (PDS)

For preparation of PDS samples, native cysteine residues were mutated as follows: C28A, C85A, C105L, C141A. The resulting Csx23 AALA mutant was used for site-specific mutagenesis and site-directed spin labelling, yielding the three Csx23 AALA constructs V52C, N59C, and N62C, with the introduced cysteines located in the transmembrane region. All mutants were tested for activity by the plasmid challenge assay described above and purified in the same manner as wild type Csx23. Protein samples were reduced with DTT (5 mM) for 2 h at 4 °C. DTT was removed by passing the sample through a PD MiniTrap G-25 column (Cytiva) pre-equilibrated with 20 mM Tris, 200 mM NaCl, pH 8.0, 0.06% DDM in D_2_O. Csx23 variants were labelled with a 10-fold molar excess of MTSL ((1-oxyl-2,2,5,5-tetramethyl-3-pyrroline-3-methyl)methanethiosulfonate; Santa Cruz Biotechnology) overnight at 4 °C. The label was removed in the same manner as DTT. MTSL-labelled protein was concentrated (Vivaspin® 500 centrifugal filter, 10 kDa MWCO PES membrane, Sartorius). Labelling efficiencies for each Csx23 mutant were assessed by continuous wave (CW) EPR measurements.

For reconstitution into nanodiscs, deuterated 20 mM bisTris, 200 mM NaCl, pH 7.0 (dEPR-ND buffer) was used throughout. DMPC (1,2-myristoyl-*sn*-glycero-3-phosphocholine, Avanti Lipids, Inc.) was dried under nitrogen to a thin film from a chloroform solution. Residual solvent was removed by high vacuum. The DMPC film was resuspended in dEPR-ND buffer to 4 mg ml^-1^ and sonicated for 5 min. The membrane scaffold protein MSP1D1 [52] was used as a scaffolding protein as described previously [53]. Briefly, MTSL-labelled Csx23 (3.6 µM), MSP1D1, and DMPC were mixed at a molar ratio of 1 : 2 : 130 in 1 ml volume in dEPR-ND buffer and Triton X-100 was added to 1.5 mM final concentration. The mixture was allowed to stand at room temperature in the dark for 30 min before the detergent was removed by successive addition of BioBeads SM-2 (BioRad). Nanodiscs were concentrated using Vivaspin® 500 centrifugal filters (10 kDa MWCO PES membrane).

PDS samples were prepared at a final monomer concentration of 50 – 75 µM for micellar Csx23, or 80 – 100 µM for Csx23 reconstituted into nanodiscs, in the absence or presence of a two-fold molar excess of cA_4_ (30 minute incubation time). 50% (v/v) deuterated glycerol (Deutero) was used for cryoprotection. The samples with a final volume of 65 µL were transferred to 3 mm quartz EPR tubes which were immediately frozen in liquid nitrogen.

### Room-temperature CW EPR

Room-temperature CW EPR measurements to assess labelling efficiency were performed using a Bruker EMX 10/12 spectrometer equipped with an ELEXSYS Super Hi-Q resonator at an operating frequency of ∼9.9 GHz (X-band) with 100 kHz modulation. Samples were recorded using a 120 G field sweep centred at 3505 G, a time constant of 20.48 ms, a conversion time of 18.67 ms, and 1714 points resolution. An attenuation of 20.0 dB (2 mW power) and a modulation amplitude of 0.7 G were used. Csx23 samples were measured in 20 μL capillaries at ∼30 μΜ monomer concentration and double integrals were compared to 4-hydroxy-TEMPO (4-hydroxy-2,2,6,6-tetramethylpiperidine 1-oxyl; Acros) as a standard. Labelling efficiency was ∼63% for the N62R1 mutant and ∼73% for both V52R1 and N59R1 mutants; samples showed negligible free spin label contribution and the shape of the spectra suggested low mobility of the label (Supplementary Figure S9).

### Pulse dipolar EPR spectroscopy (PDS)

PDS experiments were performed on a Bruker ELEXSYS E580 spectrometer with an overcoupled 3 mm cylindrical resonator (ER 5106QT-2w), operating at Q-band frequency (34 GHz). Pulses were amplified by a pulse travelling wave tube (TWT) amplifier (Applied Systems Engineering) with nominal output of 150 W. Temperature was controlled using a cryogen-free variable temperature cryostat (Cryogenic Ltd) operating in the 3.5 to 300 K temperature range.

Pulse electron-electron double resonance (PELDOR) experiments were performed at 50 K with the 4-pulse DEER [54, 55] pulse sequence (π/2(V_A_) – τ_1_ – π(V_A_) – (τ_1_ + *t*) – π(V_B_) – (τ_2_ - t) – π(V_A_) – τ_2_ – echo) as described previously [56], with a frequency offset (pump – detection frequency) of +80 MHz (∼3 mT). Shot repetition times (SRT) were set to 1.5 ms; τ_1_ was set to 380 ns, and τ_2_ was set to 4000 ns for the samples in detergent and to 2700 ns for those reconstituted into nanodiscs. The echo decays as function of available dipolar evolution time were assessed from refocused echo decays by incrementing τ_2_ in the 4 pulse DEER sequence from a start value of 760 ns and omitting the V_B_ inversion pulse. Pulse lengths were 16 and 32 ns for ν/2 and ν detection.

Measurements were performed with a reduced inversion efficiency of the pump pulse by approximately 50% to minimise multispin effects, using rectangular pulses from an arbitrary waveform generator (AWG, Bruker) with a 12 ns ELDOR pump pulse width and a 16-step phase cycle [57–60]. The pump pulse was placed on the resonance frequency of the resonator and applied to the maximum of the nitroxide field-swept spectrum. An 8-step nuclear modulation averaging with 16 ns increments was used for all experiments. Experiments ran for typically 3 – 4 h (micellar Csx23 samples) or 24 h (nanodisc samples).

PDS experiments were analyzed using DeerAnalysis2022 [61]. PDS data were first background-corrected using a 3-dimensional homogeneous background function and ghost suppression (power-scaling) for a four-spin system [62], before Tikhonov regularization followed by statistical analysis using the validation tool in DeerAnalysis2022, varying background start from 5 to 80% of the trace length in 16 trials. Resulting background start time for the best fit was then used as starting point for a second round of Tikhonov regularization followed by a second round of statistical analysis, this time including the addition of 50% random noise in 50 trials, resulting in a total of 800 trials. The regularization parameter α was chosen according to the GCV [63] or L-curve corner criterion [64] and the goodness-of-fit.

For comparison, raw PDS data were subjected to the ComparativeDEERAnalyzer (CDA) version 2.0 within DeerAnalysis2022 (DEERNet Spinach SVN Rev 5662 [65] and DeerLab 0.9.1 [66] Tikhonov regularization) for user-independent data processing and analysis, in line with current recommendations [67]. CDA reports are provided as shown in Table S2. The EPR research data underpinning this publication can be accessed at https://doi.org/.

### Modelling for PDS measurements

Distance distributions were modelled based on the AF2 structure obtained for the Csx23 full-length tetramer (PDB rank_1_model_3_ptm_seed_0_relaxed provided in the underpinning data). R1 moieties were introduced at residues 52, 59, or 62 of all four chains of the tetramer using both mtsslWizard [43] within the mtsslSuite server-based modelling software [68] and “tight” settings, and Multiscale Modeling of Macromolecules (MMM) [42] using “ambient temperature” settings. Cartoon structural representations of spin-labelled Csx23 constructs were generated using Pymol (Schrödinger Inc.).

### Crystallisation of Csx23

Csx23 tetramer at 10 mg mL^−1^ was mixed in a 1:2 molar ratio with cA_4_ and incubated at room temperature for 30 min, before centrifugation at 13,000 rpm prior to crystallisation. Sitting drop vapour diffusion experiments were set up at the nanoliter scale using commercially available crystallisation screens and incubated at 293 K. Crystals used for data collection were evident after 2 days and grew from a reservoir solution of 42.5% PEG 400, 0.2M LiSO_4_, 0.1 M sodium acetate pH 5, which also acted as an intrinsic cryoprotectant. As such crystals were harvested and immediately cryo-cooled prior to data collection.

### X-ray data processing, structure solution, and refinement

X-ray data were collected at a wavelength of 0.9537 Å, on beamline I04 at Diamond Light Source, at 100 K to 1.76 Å resolution. Data were automatically processed using autoPROC [69] and STARANISO [70]. The data were phased by PhaserMR [71] in the CCP4 suite [72] using a model generated by AlphaFold2 [30] implemented in Colab, with initial B-factors modelled in Phenix [73]. Model refinement was achieved by iterative cycles of REFMAC5 [74] with manual model manipulation in COOT [75]. Electron density for cA_4_ was clearly visible in the maximum likelihood/σA weighted *F*_obs_-*F*_calc_ electron density map at 3σ. The coordinates for cA_4_ were generated in ChemDraw (Perkin Elmer) and the library was generated using acedrg [76], before fitting of the molecule in COOT. Model quality was monitored throughout using Molprobity [77] (score 1.48; centile 94). Ramachandran statistics are 96.3% favoured, 0% disallowed. Data and refinement statistics are shown in Table S3. The coordinates and data have been deposited in the Protein Data Bank with deposition code 8QJK.

## Acknowledgements

Dr Laura Remmel is thanked for purification of MSP1D1, and Dr Samantha Pitt for helpful discussions. This work was supported by a grant from the Biotechnology and Biological Sciences Research Council (Grant REF BB/T004789/1 to MFW and TMG) and the Engineering and Physical Sciences Research Council (Grant REF BB/T004789/1 to KA, BEB and MFW). EPR equipment was funded by BBSRC (BB/R013780/1 and BB/T017740/1).

## Author contributions

Sabine Grüschow, Investigation, Methodology, Formal analysis, Visualisation, Writing-original draft preparation, Writing-review and editing; Stuart McQuarrie, Investigation, Methodology, Formal Analysis, Writing-review and editing; Stephen McMahon, Investigation, Methodology, Formal Analysis, Writing-review and editing; Katrin Ackermann, Investigation, Methodology, Formal Analysis, Writing-review and editing; Bela Bode, Investigation, Methodology, Formal Analysis, Writing-review and editing; Tracey M. Gloster, Funding acquisition, Supervision, Visualisation, Writing-review and editing; Malcolm F. White, Conceptualisation, Formal analysis, Supervision, Project administration, Funding acquisition, Writing-original draft preparation, Writing-review and editing.

## Supporting Information

**Figure S1.**
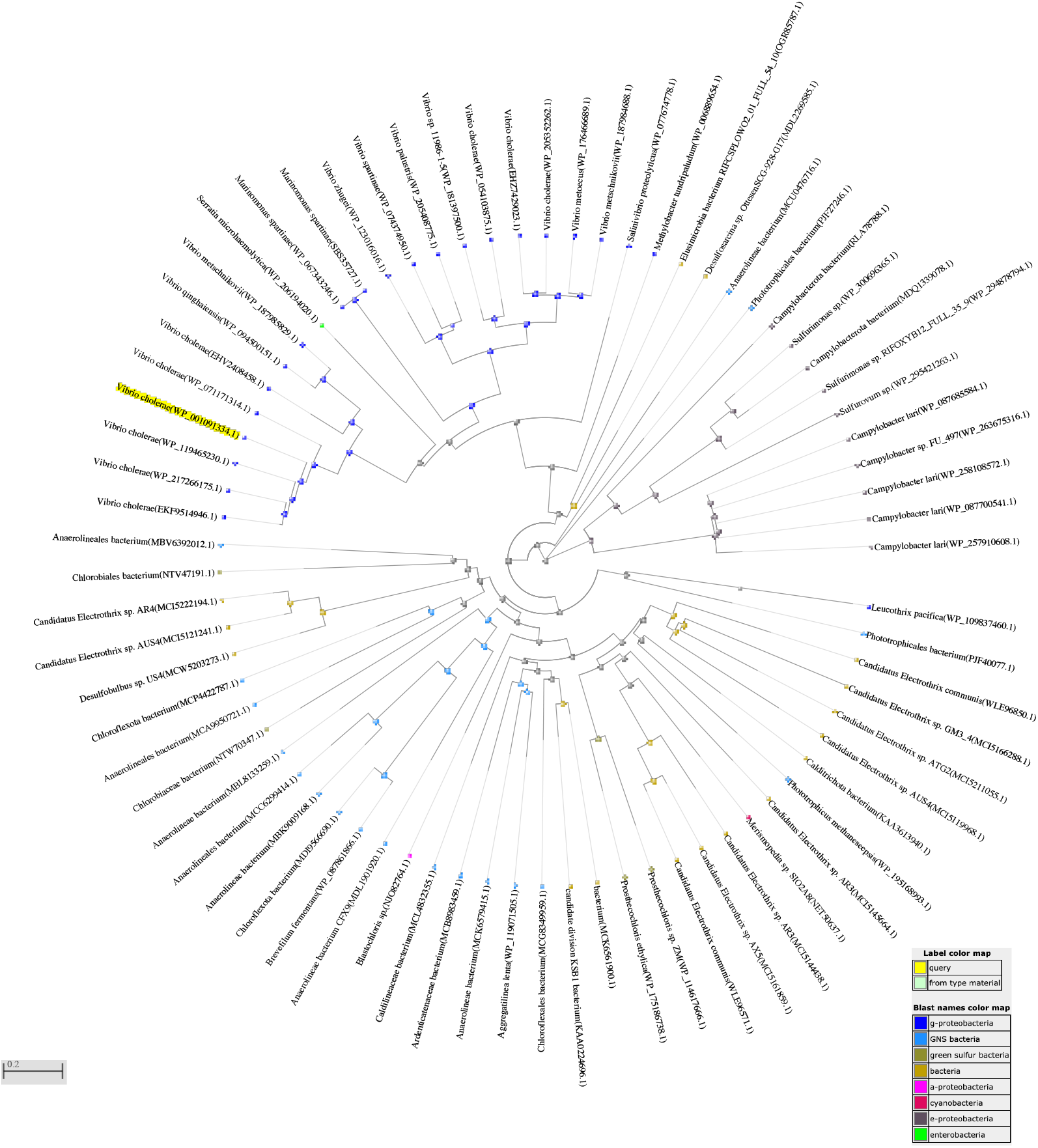
Distance tree of PSI-BLAST results using Csx23 (WP_001091334.1, highlighted) as query (NCBI website). The tree is based on pairwise alignments with fast minimum evolution (Grishin distance model, 0.85 maximum sequence difference) as method for tree construction.

**Figure S2.**
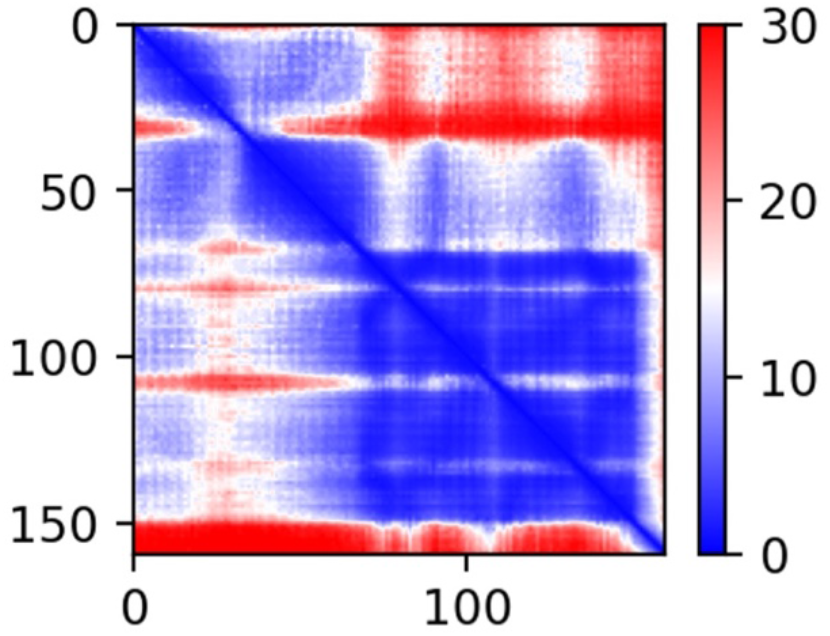
Predicted Aligned Error (PAE) plot output from AF2 for FL Csx23. The colour at (x, y) indicates the expected position error at residue x (x axis) if the predicted and true structures were aligned on residue y (y axis). Coloured from blue (low error) to red (high error) as indicated on side bar in Å. This plot shows the residue positional predictions within each domain have low error, but predictions of the residue positions between the two domains is higher.

**Figure S3.**
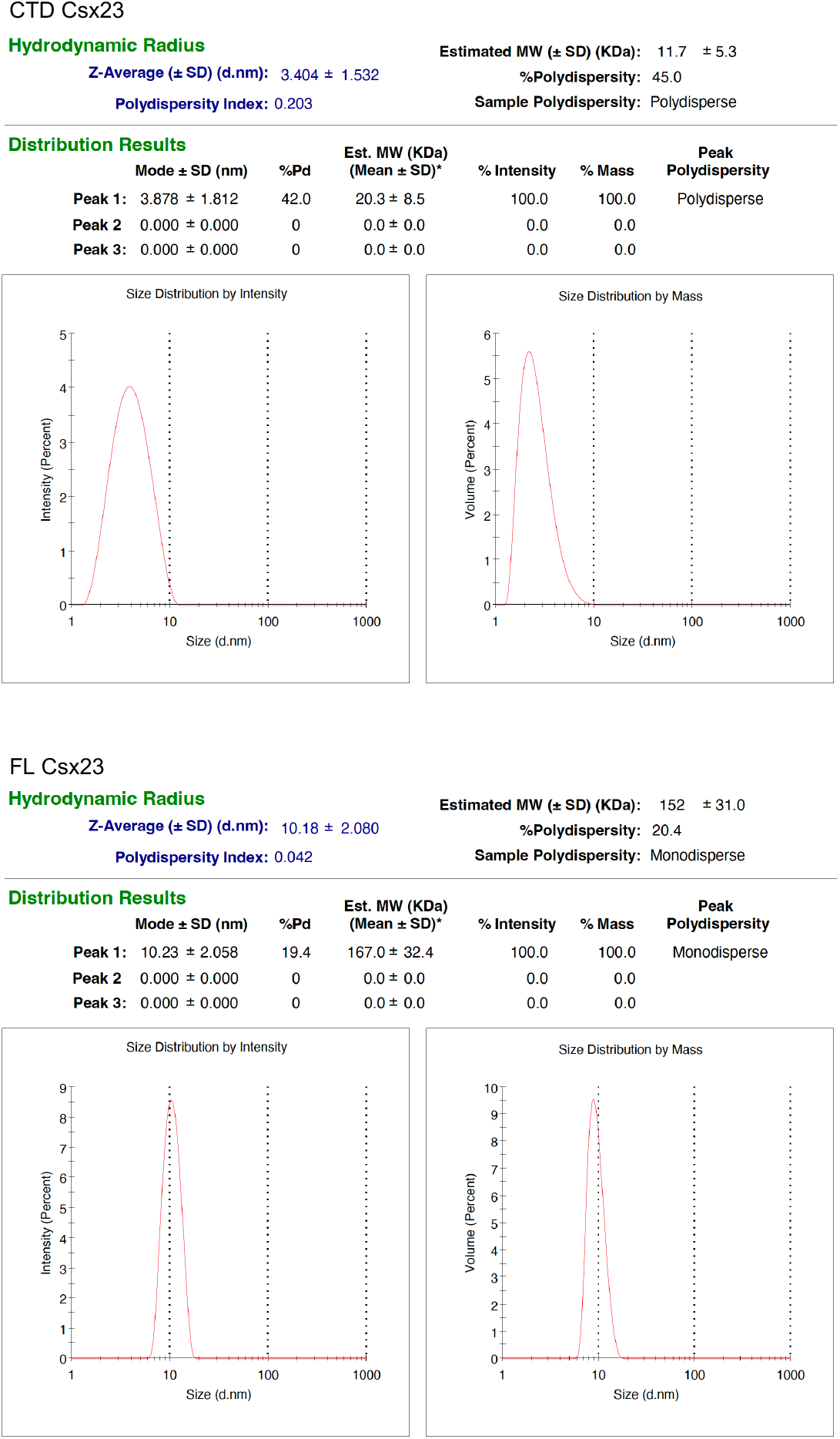
Dynamic light scattering (DLS) analysis of Csx23. The molecular weight (MW) was calculated from the hydrodynamic radius using the Zetasizer software assuming a globular shape. The soluble, C-terminal domain (CTD, top) of Csx23 was measured in the absence of detergent; the size measured by DLS fits well with the CTD being monomeric (expected MW 10.4 kDa). Full length (FL, bottom) Csx23 was measured in the presence of 0.1% DDM; the size measured for the full length detergent:protein complex is at least 10 times larger than that obtained for the CTD. This may suggest an oligomeric form for FL Csx23, however, the presence of detergent precludes further interpretation.

**Figure S4.**
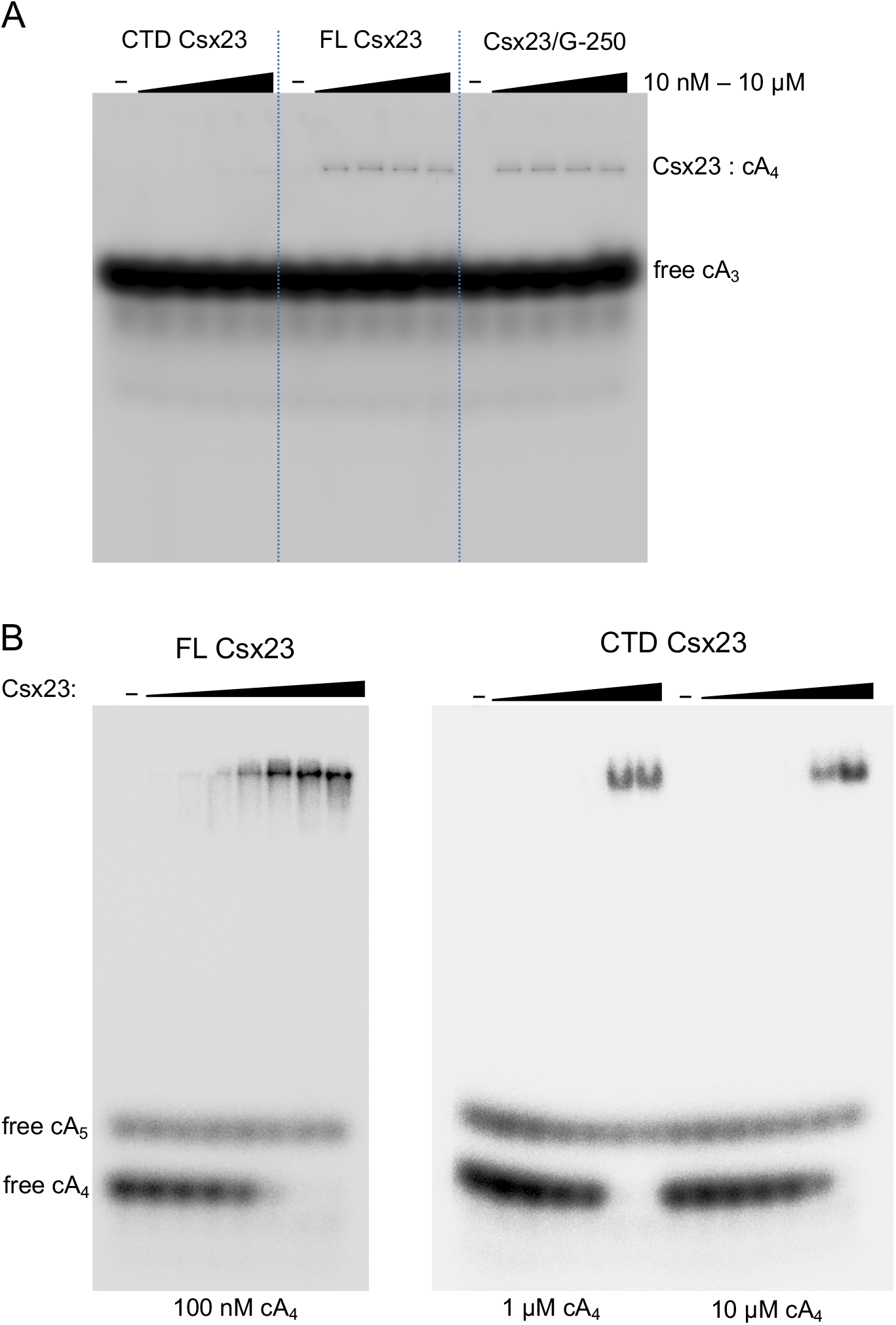
Uncropped gels for Csx23 : cOA binding assays. **A:** EMSA with radiolabelled VmeCmr-derived cA_3_ and FL Csx23 or Csx23 CTD. A 10-fold serial dilution of Csx23 was incubated with approximately 1.5 µM [^32^P]-cOA each. Csx23/G-250: Coomassie G-250 was added prior to loading. **B:** Full images for Figure 2. EMSA with radiolabelled SsoCsm-derived cA_4_. Protein concentrations were 25, 50, 100, 200, 400, 800 and 1600 nM for FL Csx23 and 0.4, 1, 4, 10, 40 and 100 µM for Csx23 CTD.

**Figure S5.**
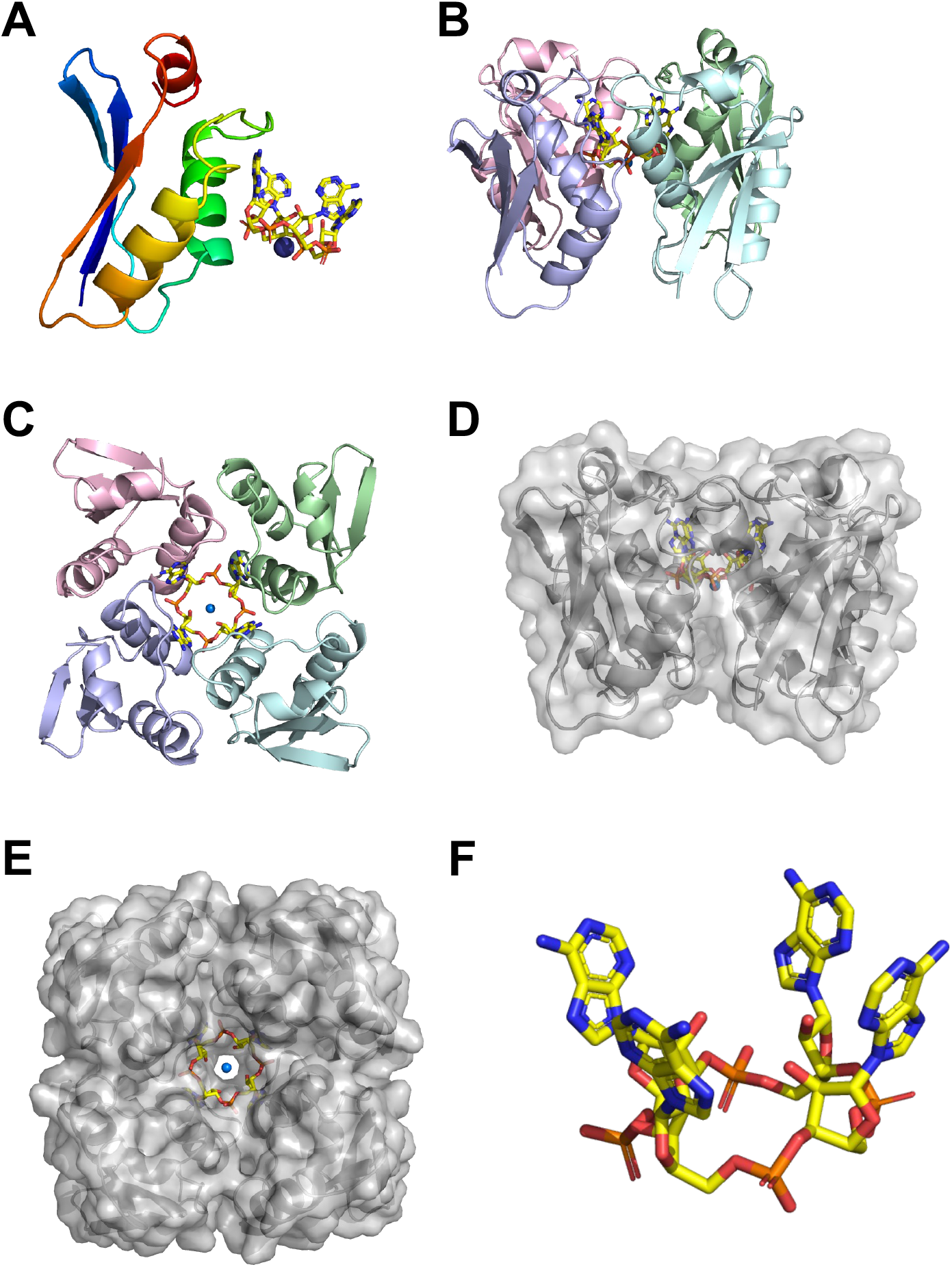
Structure of Csx23 CTD. **A**: Cartoon representation of monomeric Csx23 CTD coloured from N-(blue) to C-terminus (red). cA_4_ is shown in sticks (coloured by element with carbon in yellow) and the sodium ion as a blue sphere for orientation. **B**: ‘Side’ view and **C**: ‘top’ view of tetrameric Csx23 CTD, shown in cartoon representation with each monomer in a different colour. cA_4_ is shown in sticks (coloured by element with carbon in yellow) and the sodium ion as a blue sphere. **D**: ‘Side’ view and **E**: ‘top’ view of tetrameric Csx23 CTD shown in surface representation. Colouring as in panels **B**/**C**. **F**: Stick representation of cA_4_, coloured by element with carbon in yellow.

**Figure S6.**
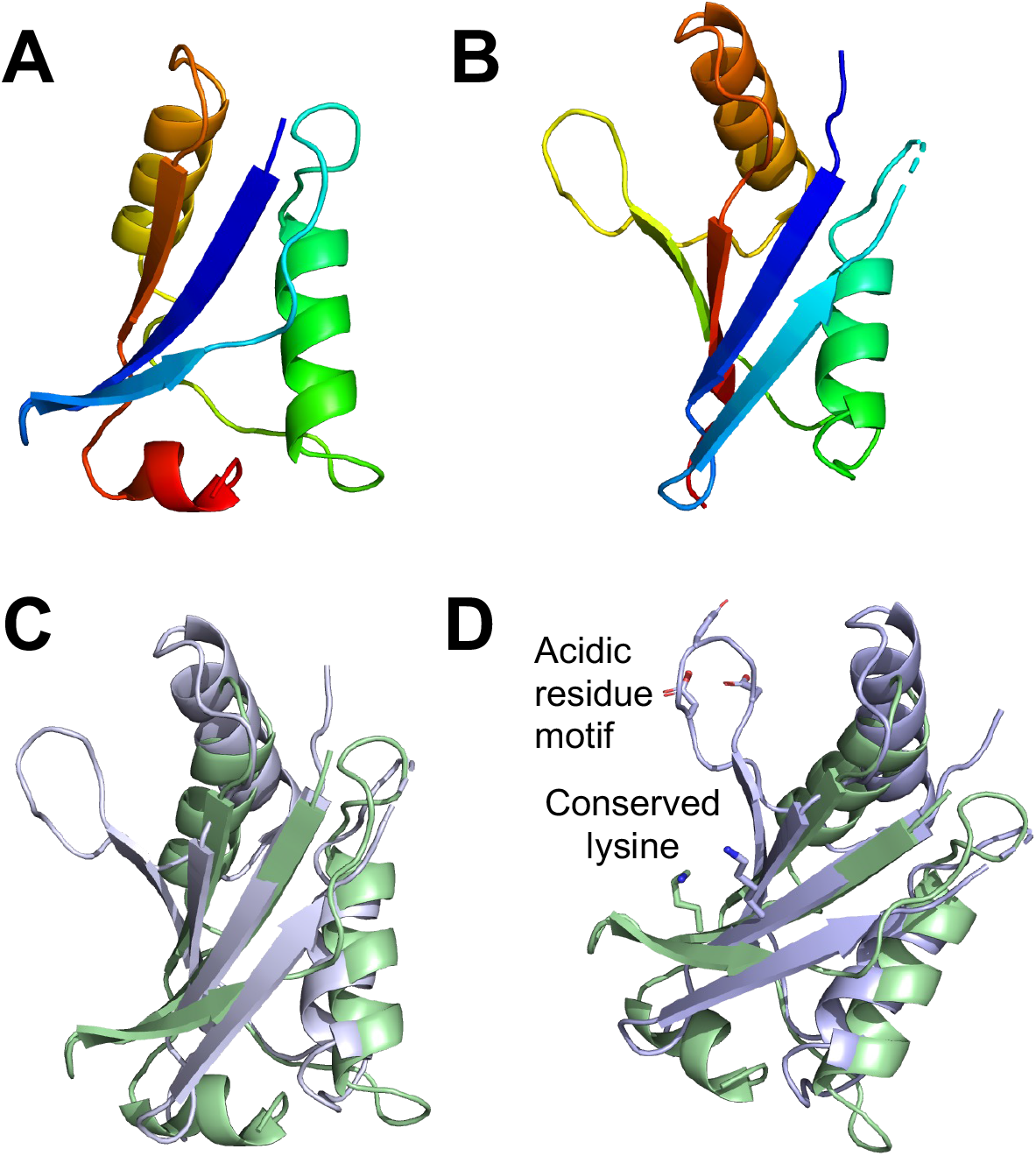
Comparison of the structures of Csx23 CTD and the PB1 domain from protein kinase C zeta type. **A**: Cartoon representation of monomeric Csx23 CTD coloured from N-(blue) to C-terminus (red). **B**: Cartoon representation of the PB1 domain from protein kinase C zeta type from rat (PDB: 4MJS) coloured from N-(blue) to C-terminus (red). **C**: Superimposition of Csx23 CTD (green cartoon) and the PB1 domain from protein kinase C zeta type (light blue cartoon). **D**: Superimposition of Csx23 CTD (green cartoon) and the PB1 domain from protein kinase C zeta type (light blue cartoon), highlighting a conserved lysine residue at the end of the first beta-strand in each protein, and the additional region containing acidic residues present only in the PB1 domain.

**Figure S7.**
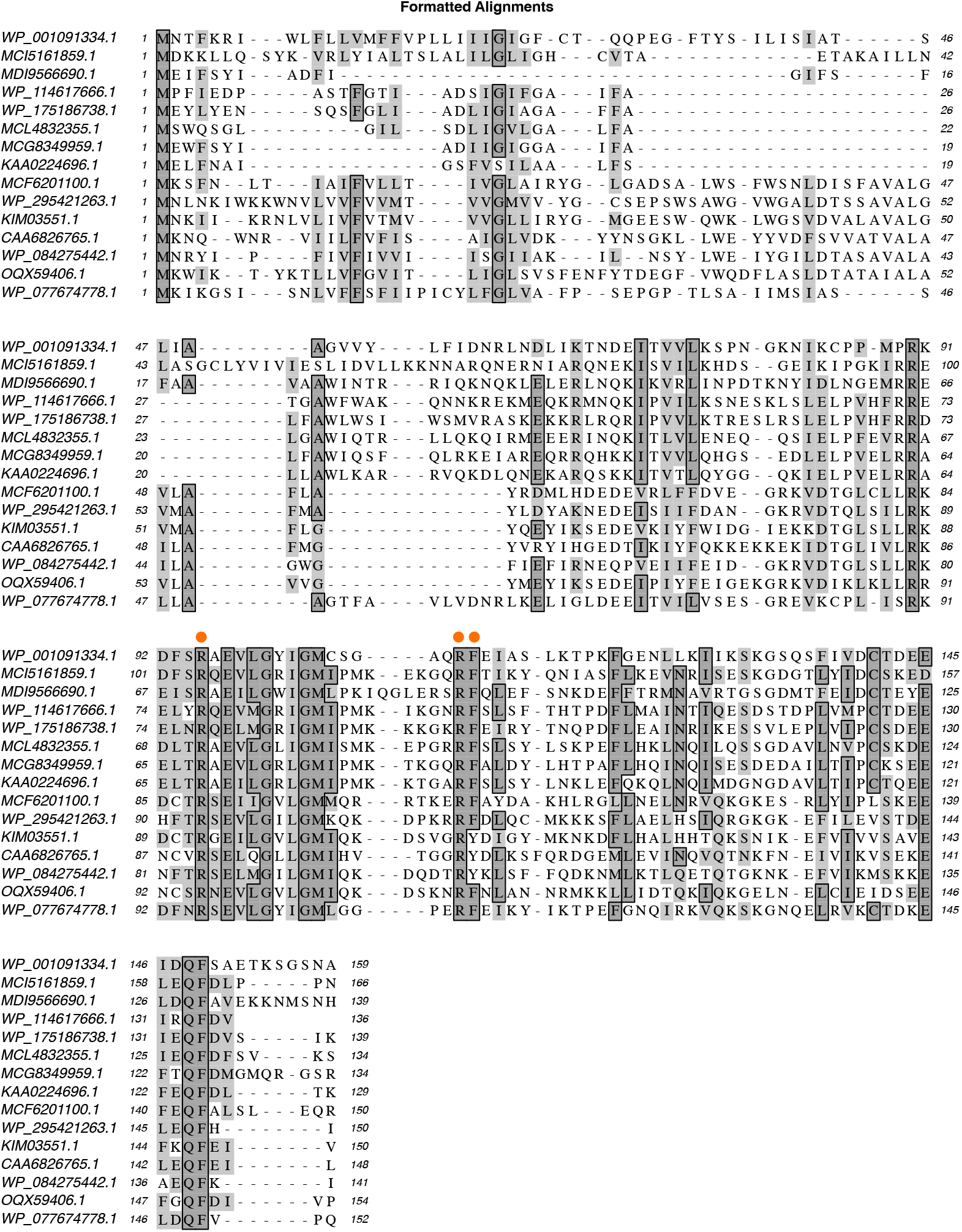
Multiple sequence alignment of *V. cholerae* Csx23 (WP_001091334.1) and diverse homologues, aligned using MUSCLE. Residues targeted by mutagenesis (R95, R110, and F111) are marked by a dot.

**Figure S8.**
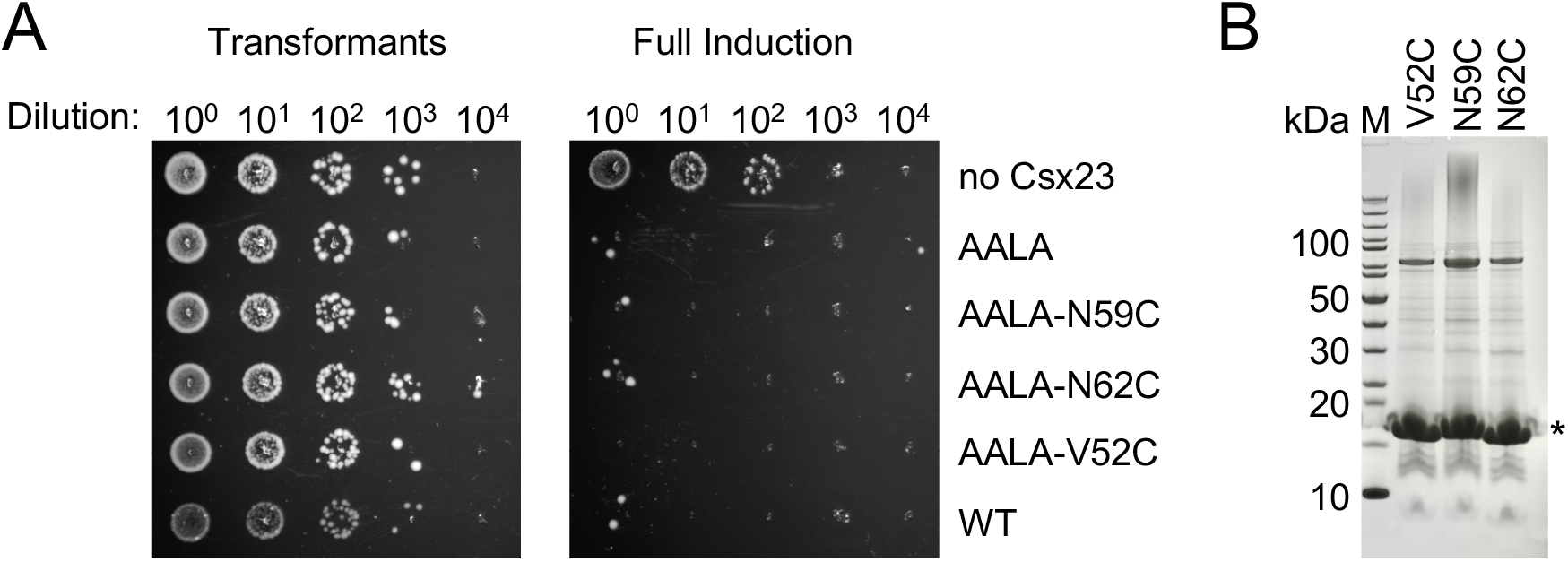
Characterisation of Csx23 variants used for spin-labeling and EPR experiments. **A:** Plasmid challenge assay, demonstrating that all variants retained biological function. Figure representative of two technical replicates. **B:** SDS-PAGE of purified variants of Csx23.

**Figure S9.**
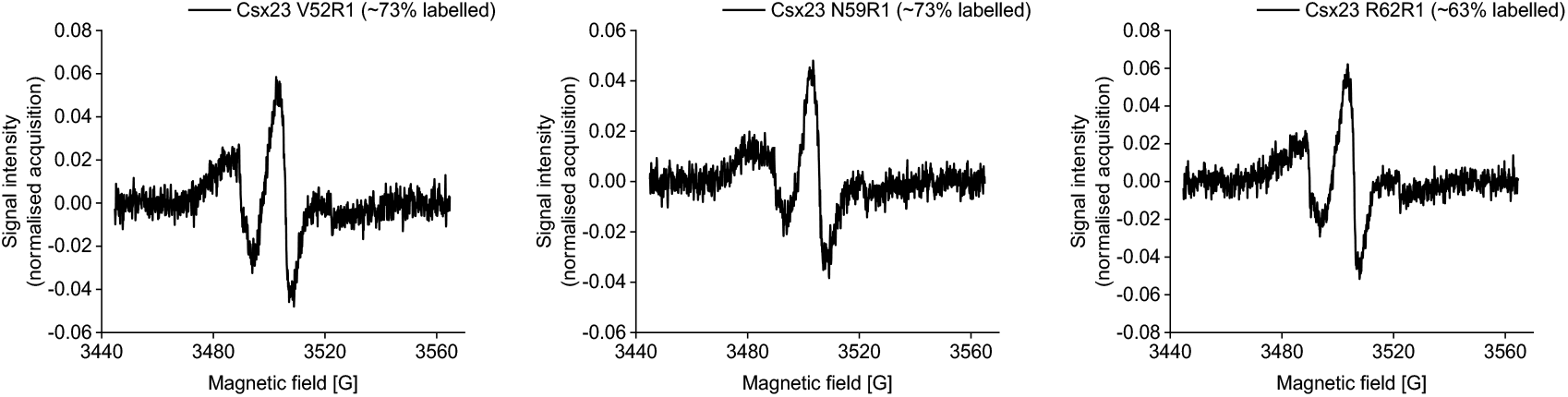
Continuous wave (CW) EPR spectra. Individual CW EPR spectra for MTSL-labelled Csx23 variants as indicated. Details of the constructs and corresponding labelling efficiencies are given on each plot.

**Figure S10.**
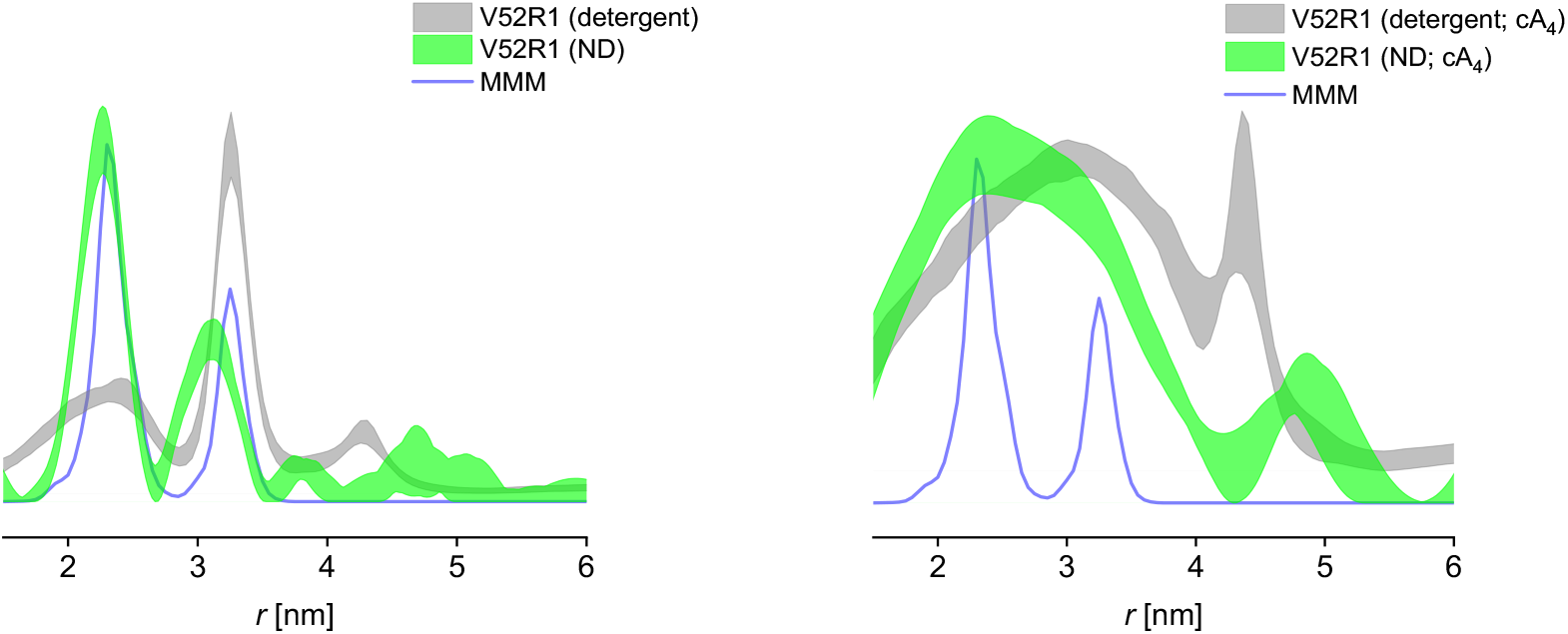
PELDOR data for the Csx23 AALA V52R1 mutant in absence (left) and presence (right) of cyclic nucleotide (cA_4_), comparing micellar protein (detergent) to protein reconstituted in nanodiscs (ND). The predicted distribution based on the AF2 tetramer predicted structure of Csx23 is shown in blue.

**Figure S11.**
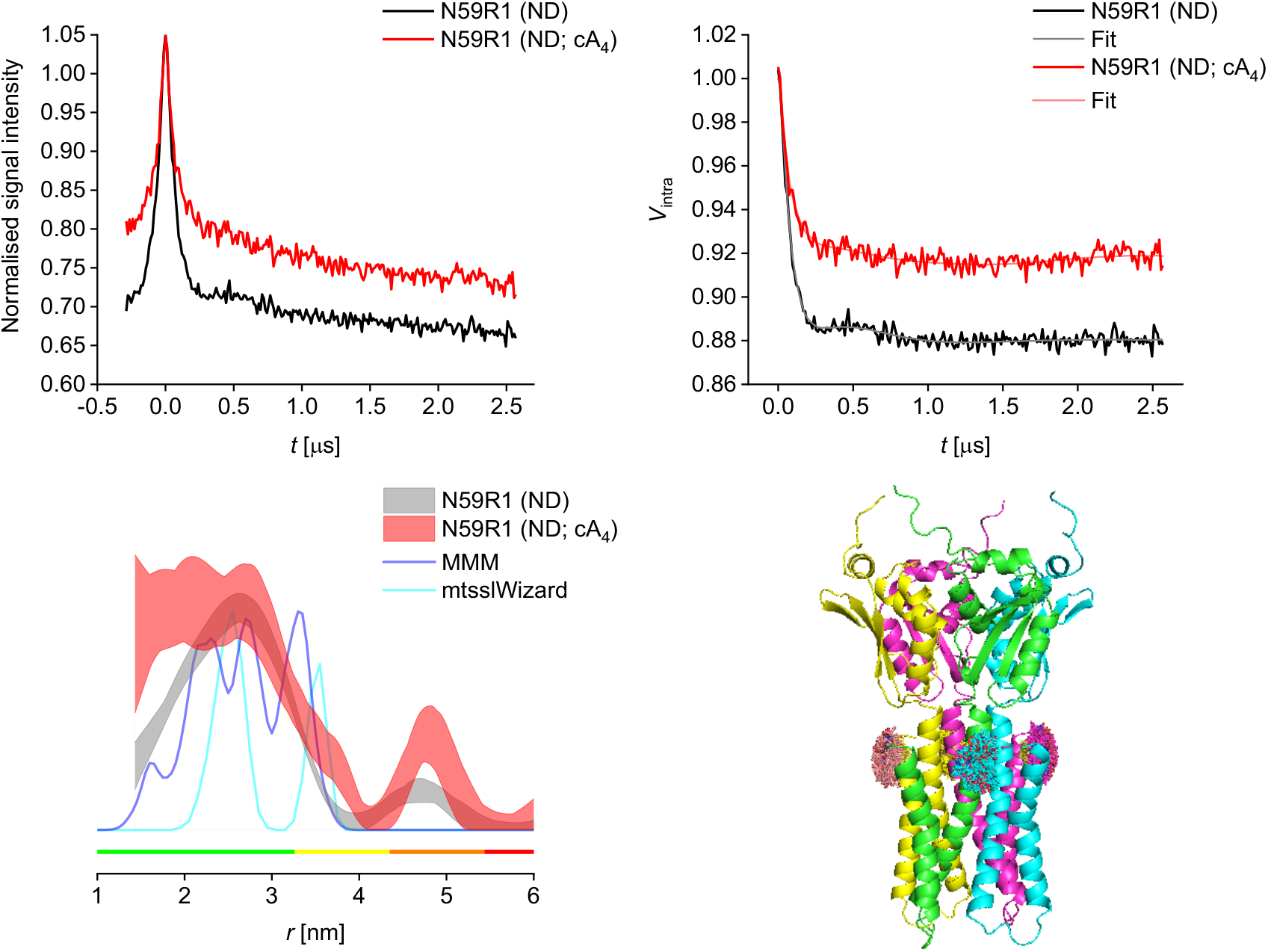
PELDOR data for the Csx23 AALA N59R1 mutant reconstituted in nanodiscs (ND) in presence (red) and absence (grey) of cyclic nucleotide (cA_4_). Raw PELDOR data (top left) and background-corrected traces with fits (top right); overlay of corresponding distance distributions shown as 95% confidence bands with predicted distributions from MMM and mtsslWizard based on AF2 predicted structure (bottom left), colour bars indicate reliability ranges (green: shape reliable; yellow: mean and width reliable; orange: mean reliable; red: no quantification possible); cartoon representation of AF2 predicted tetrameric structure of the spin-labelled Csx23 AALA N59R1 tetramer (bottom right).

**Figure S12.**
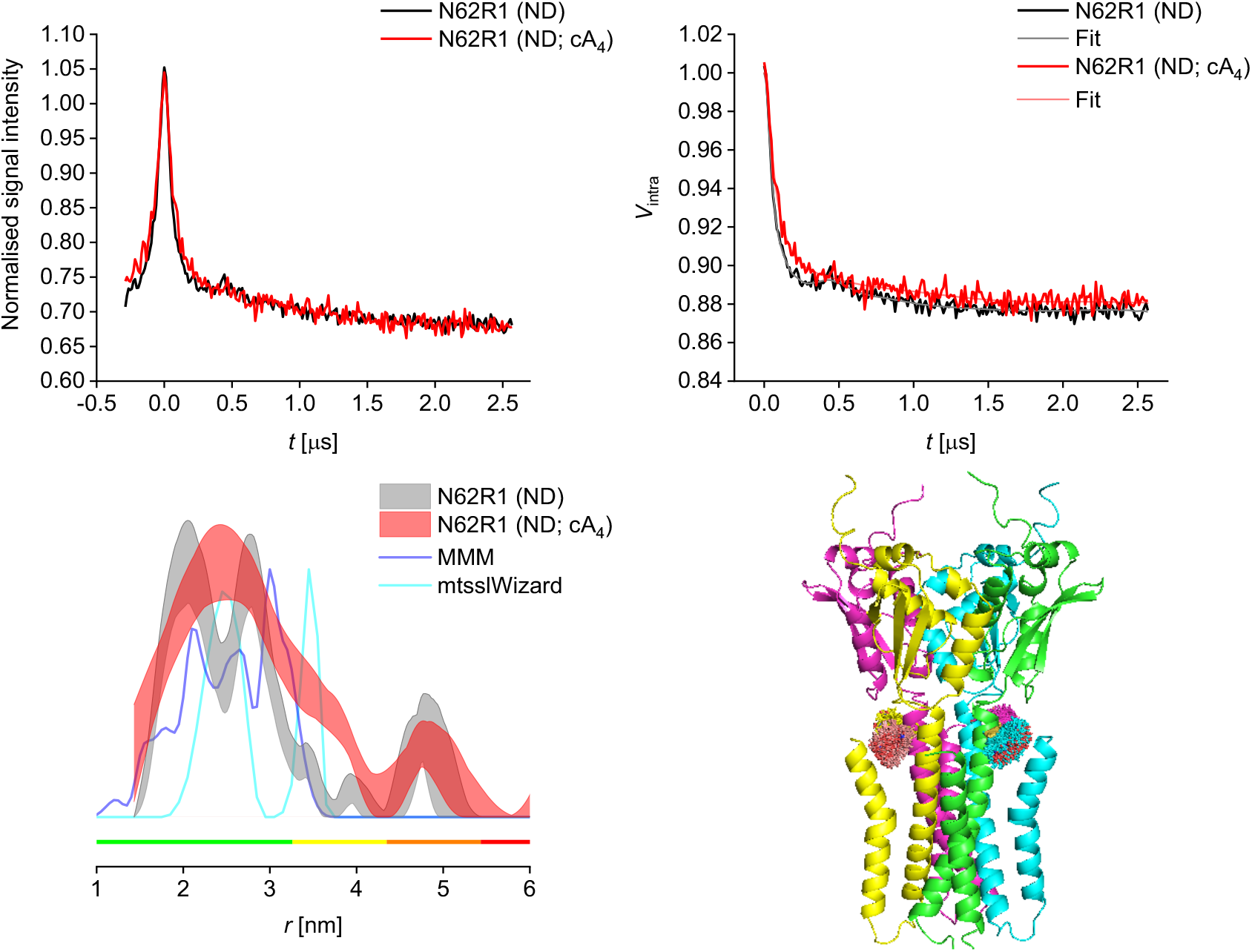
PELDOR data for the Csx23 AALA N62R1 mutant reconstituted in nanodiscs (ND) in presence (red) and absence (grey) of cyclic nucleotide (cA_4_). Raw PELDOR data (top left) and background-corrected traces with fits (top right); overlay of corresponding distance distributions shown as 95% confidence bands with predicted distributions from MMM and mtsslWizard based on AF2 structure (bottom left), colour bars indicate reliability ranges (green: shape reliable; yellow: mean and width reliable; orange: mean reliable; red: no quantification possible); cartoon representation of AF2 predicted tetrameric structure of the spin-labelled Csx23 AALA N62R1 tetramer (bottom right).

**Figure S13.**
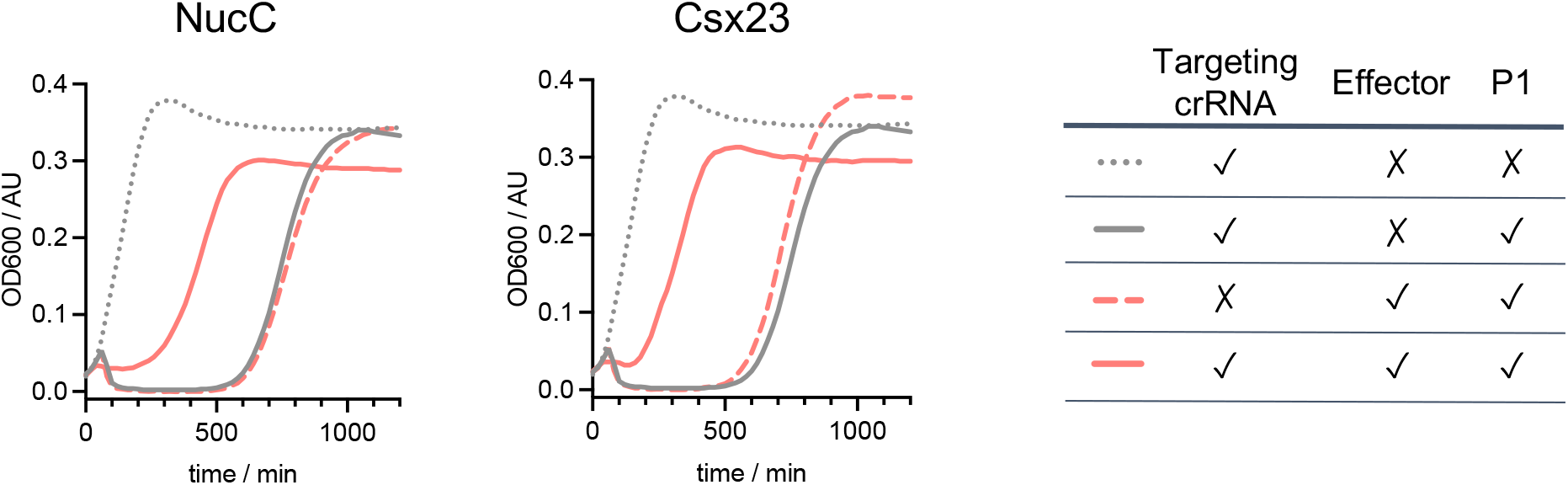
Phage P1 immunity assay using the VmeCmr / effector system. The growth of cells harbouring either Csx23 or NucC in the absence of targeting crRNA (dashed, salmon line) is the same as that of cells not carrying an effector (solid, grey line). The combination of targeting crRNA and effector results (solid, salmon line) in significantly faster recovery of the culture at high MOIs (MOI 15).

**Table S1.**
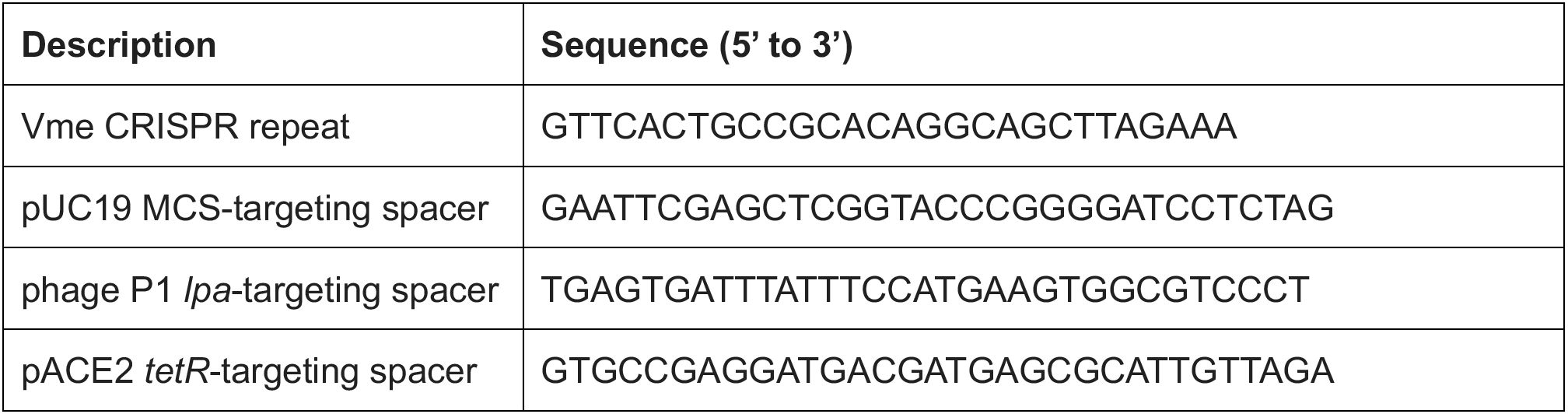
VmeCRISPR repeat and spacer sequences used in this study.

**Table S2.** Key to Comparative DeerAnalyzer (CDA2.0) Reports. ND = reconstituted into nanodiscs.

| **CDA report** | **Csx23 sample** |
| --- | --- |
| 220913_KAq200.16_DEER_128032_pi_half_comparative_DEER_analyzer_report | Csx23 AALA V52R1 |
| 220913_KAq200.19_DEER_128032_pi_half_comparative_DEER_analyzer_report | Csx23 AALA V52R1 + cA<sub>4</sub> |
| 230210_KAq204.3_PELDOR_128032_pi_half_comparative_DEER_analyzer_report | Csx23 AALA V52R1, ND |
| 230211_KAq204.6_PELDOR_128032_pi_half_comparative_DEER_analyzer_report | Csx23 AALA V52R1 + cA<sub>4</sub>, ND |
| 230309_KAq210.3_deer_128032_pi_half_comparative_DEER_analyzer_report | Csx23 AALA N59R1, ND |
| 230312_KAq210.12_DEER_128032_pi_half_comparative_DEER_analyzer_report | Csx23 AALA N59R1 + cA<sub>4</sub>, ND |
| 230310_KAq210.6_DEER_128032_pi_half_comparative_DEER_analyzer_report | Csx23 AALA N62R1, ND |
| 230311_KAq210.9_DEER_128032_pi_half_comparative_DEER_analyzer_report | Csx23 AALA N62R1 + cA<sub>4</sub>, ND |

**Table S3.**
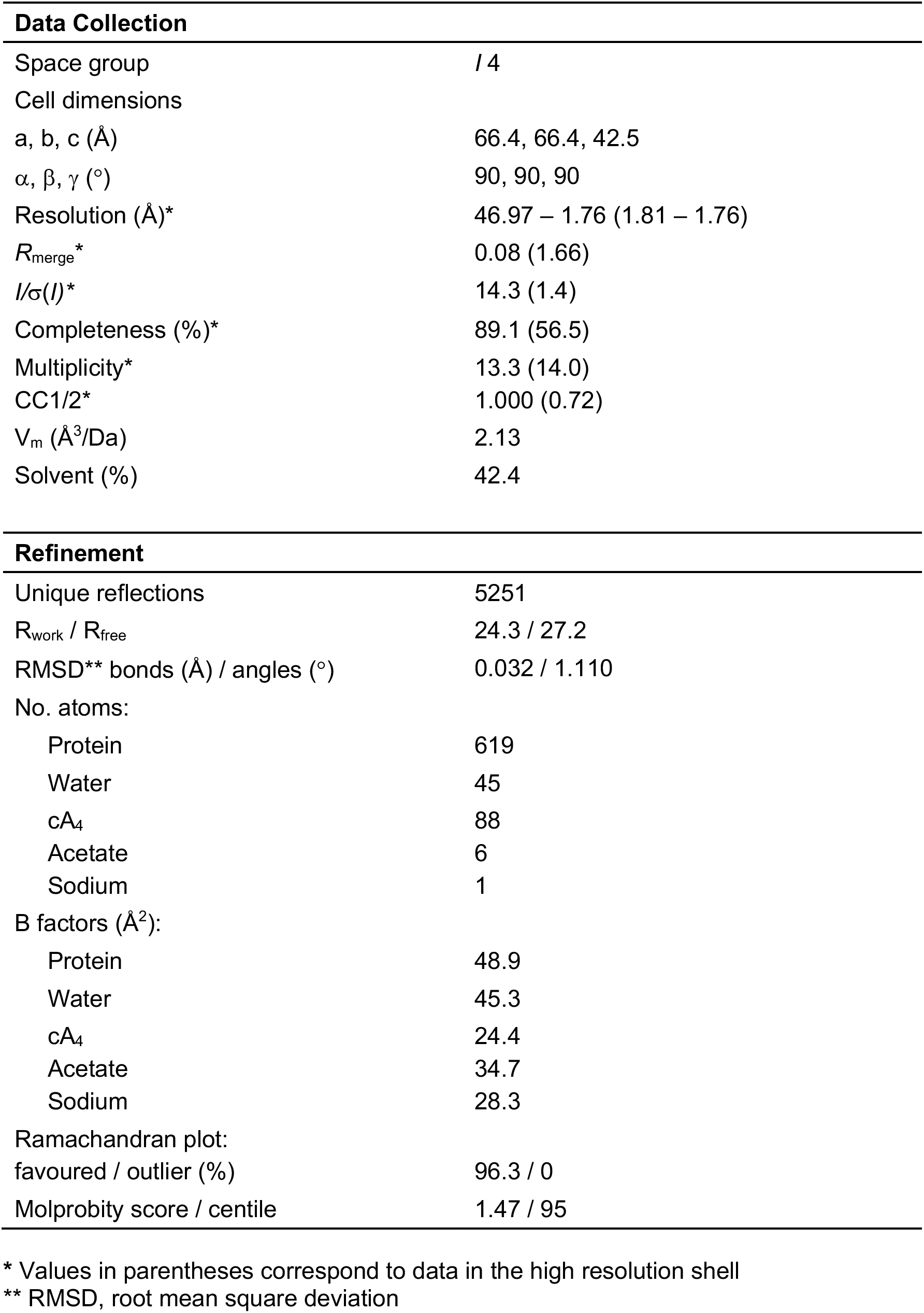
Data collection and refinement statistics for the structure of Csx23 CTD in complex with cA_4_.

